# On-demand surround suppression explains the spatiotemporal window of the tilt illusion

**DOI:** 10.64898/2026.10.01.755975

**Authors:** Tomoya Nakamura, Lana Okubo, Kiyofumi Miyoshi, Ikuya Murakami

## Abstract

What we see at one location depends on what surrounds it. Such contextual effects are thought to arise from neural signals that travel across the retinotopic representations. Because these signals need more time to travel longer distances, a more distant surround should influence perception most strongly when presented earlier, compensating for its delay. We tested this prediction in human observers using a visual illusion in which a central target appears tilted away from the orientation of its surround (tilt illusion). Measuring the temporal window of this illusion using psychophysical reverse correlation revealed the opposite: the most effective surround occurred progressively later as distance increased. Computational modelling demonstrated that this reversal is explained if the visual system sends out a query that recruits contextual information on demand. Perceptual appearance may thus be constructed through temporally unfolding interactions in which context is actively gathered, not passively received.

## Introduction

No part of a natural scene is statistically independent of its surroundings. Unlike white noise, natural images are highly structured: luminance, colour, and texture vary smoothly within the same object or surface, but change abruptly across the boundaries between them. The visual system is considered to exploit such statistical regularity, evaluating a local feature not in isolation but relative to the features that surround it^1–3^.

Such spatial comparison is actually implemented in the biological visual system. We focus here on the orientation domain, which has been characterized in the greatest detail, from single neuron physiology to psychophysics. Orientation-selective neurons in the primary visual cortex (V1), for example, reduce their responses to a central target grating of their preferred orientation when surrounding regions contain gratings of similar orientations, a phenomenon known as iso-orientation surround suppression^4–7^.

Contextual modulation even changes what we perceive, that is, the visual appearance of scene elements. One striking perceptual correlate of the iso-orientation surround suppression is the *tilt illusion*, the perceptual bias in which a target’s orientation is systematically biased by the orientation of its surround, typically away from the surround orientation^8,9^. Because lateral inhibition acts most strongly among neurons tuned to similar orientations, neurons preferring orientations close to that of the surround become less responsive among others. Consequently, at a population level, the target orientation is represented as repelled away from surrounding orientation^10–13^, consistent with the perceptual repulsion.

Despite their success in predicting the static or equilibrium state of the final percept, however, these normative and mechanistic accounts do not consider the neural dynamics that may arise from the time delay in integrating signals from spatially distant locations. Indeed, neural delays of iso-orientation surround suppression increase with distance from the central stimulus^14^ (see also ref.^15^). If the tilt illusion involves iso-orientation surround suppression, such distance-dependent delays must therefore be accommodated before the percept is finalized. We reasoned that a signature of how they are accommodated can be recovered by measuring the temporal window within which the surround inducer could act on the central target: moving the inducer farther from the target should systematically move that window, by an amount set by the propagation delay.

To measure the temporal window of the tilt illusion, we took advantage of the method of psychophysical reverse correlation^16–21^. In Mareschal and Clifford’s experiment^22^, observers judged whether a briefly flashed vertical target appeared to be tilted clockwise (CW) or counter-clockwise (CCW), when an abutting inducer randomly changed its orientation in each display frame. By correlating the observer’s choice with the orientation and relative time of the inducer, they reconstructed a correlogram that quantified the contribution of each inducer to the tilt illusion as a function of time relative to the target’s onset time.

The present study identified such correlograms at several target–inducer distances. To anticipate, we found that, with increasing distance, the temporal window shifted toward the future relative to the target onset time. The shift direction is inconsistent with the account that considers only distance-dependent delay from the surround to the centre^14,15,23^, which tacitly assumes that the surround incessantly sends suppressive signals inward whenever it is stimulated and the centre passively receives them. Instead of this passive account, we propose an alternative mechanistic model in which a target onset triggers active querying that propagates query signals outward. The query establishes a new communication channel, which enables reply signals from the destination to help manage the target’s property settings such as orientation. This *active* account naturally predicts the shift to the future. Our formal modelling, based on the canonical neural computation of normalization^24,25^, quantitatively reproduced the empirical correlogram. Our view is therefore that the visual system does not simply receive its context but goes out and asks for it, actively constructing what we perceive from the answer that comes back.

## Results

To derive the temporal windows of the tilt illusion at different distances, we requested each of our ten observers to perform thousands of orientation discrimination trials (Fig. 1). In each trial, a horizontal Gabor patch (target) was flashed for 33 ms at 8 degrees of visual angle (dva) below the fixation point. The target was surrounded by an annular grating (inducer) at one of the three distances (2.0, 3.5, and 5.0 dva, referred to as the near, middle, and far conditions); inducer orientation was randomly changed every 33 ms. After a 0.3 s blank period, CW– and CCW-tilted probes were presented, and the observers were asked to choose the one that better resembled the target orientation. The target orientation was exactly horizontal except in catch trials, which occupied 21% of trials, wherein the target was tilted either +10° or −10°.

**Fig. 1.**
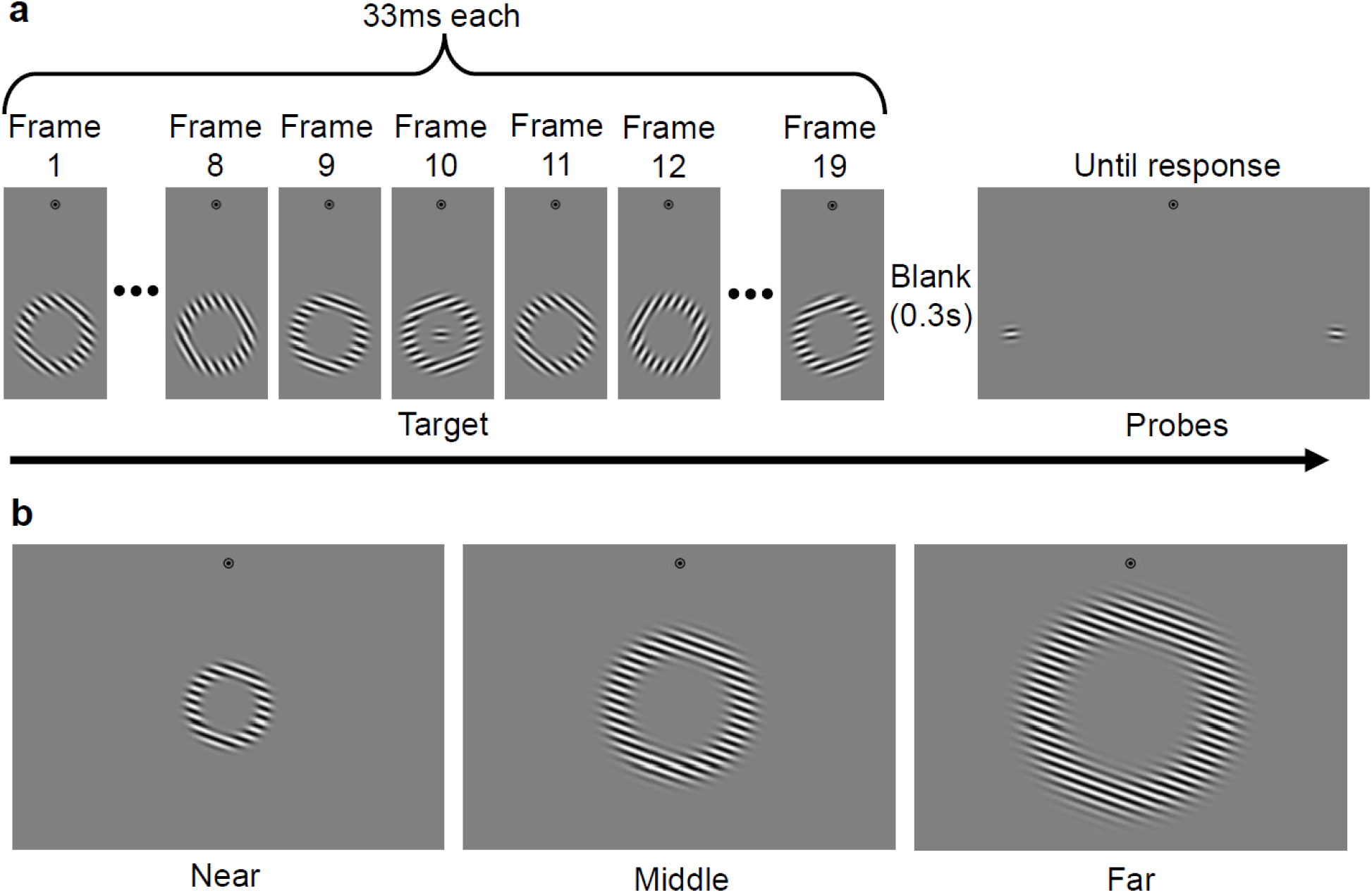
Procedure. (a) Time course of a single trial. Nineteen inducer frames were presented successively, each lasting 33 ms with a randomly chosen orientation (0°, ±20°, ±40°, ±60°, or ±80° from horizontal). A horizontal Gabor target was flashed for 33 ms at the centre of the annulus, simultaneously with the tenth inducer. After a 300-ms blank, two probes tilted +5° and −5° appeared to the left and right of the target location, and observers reported which probe better resembled the target orientation. (b) Inducers in the near (left), middle (centre), and far (right) conditions. Varying the target–inducer distance varies the cortical distance that contextual signals must travel. If this travel incurs a distance-dependent delay, the delay should manifest as a shift of the temporal window across the three conditions.

### Temporal window peaks later at greater distances

After confirming that the observers were well engaged in our task, as certified by their excellent proportions correct in the catch trials (91.4±2.5%, 96.8±1.1%, and 97.5±0.7% s.e.m. in the near, middle, and far conditions, respectively), we derived the temporal window at each distance separately. We made a correlogram to visualise the proportion of CCW judgements, for each combination of inducer orientation and inducer onset time (τ) relative to the target onset time. In this correlogram, positive deviations for the CW-inducer and negative deviations for the CCW-inducer from the chance level (50%) indicate the degree of contribution of the corresponding inducer to repulsion. Such deviations were concentrated around the moment of target presentation (τ = 0), but only when the inducer was tilted ±20° from the horizontal (Fig. 2a for a representative observer’s data; see Supplementary Fig. 1 for all observers’ data). This qualitative trend pertaining to the temporal and orientation tunings was consistent across the observers and also similar to those in Mareschal and Clifford’s study^22^. As in Mareschal and Clifford^22^, we flipped the proportion CCW for −20° inducers and merged it with the proportion CCW for +20° inducers, plotting the proportion of trials in which the target was judged as tilted opposite to the inducer (hereafter called the “proportion opposite”) against τ and fitted it with a Gaussian function (descriptive model; Fig. 2b) for each observer and condition.

**Fig. 2.**
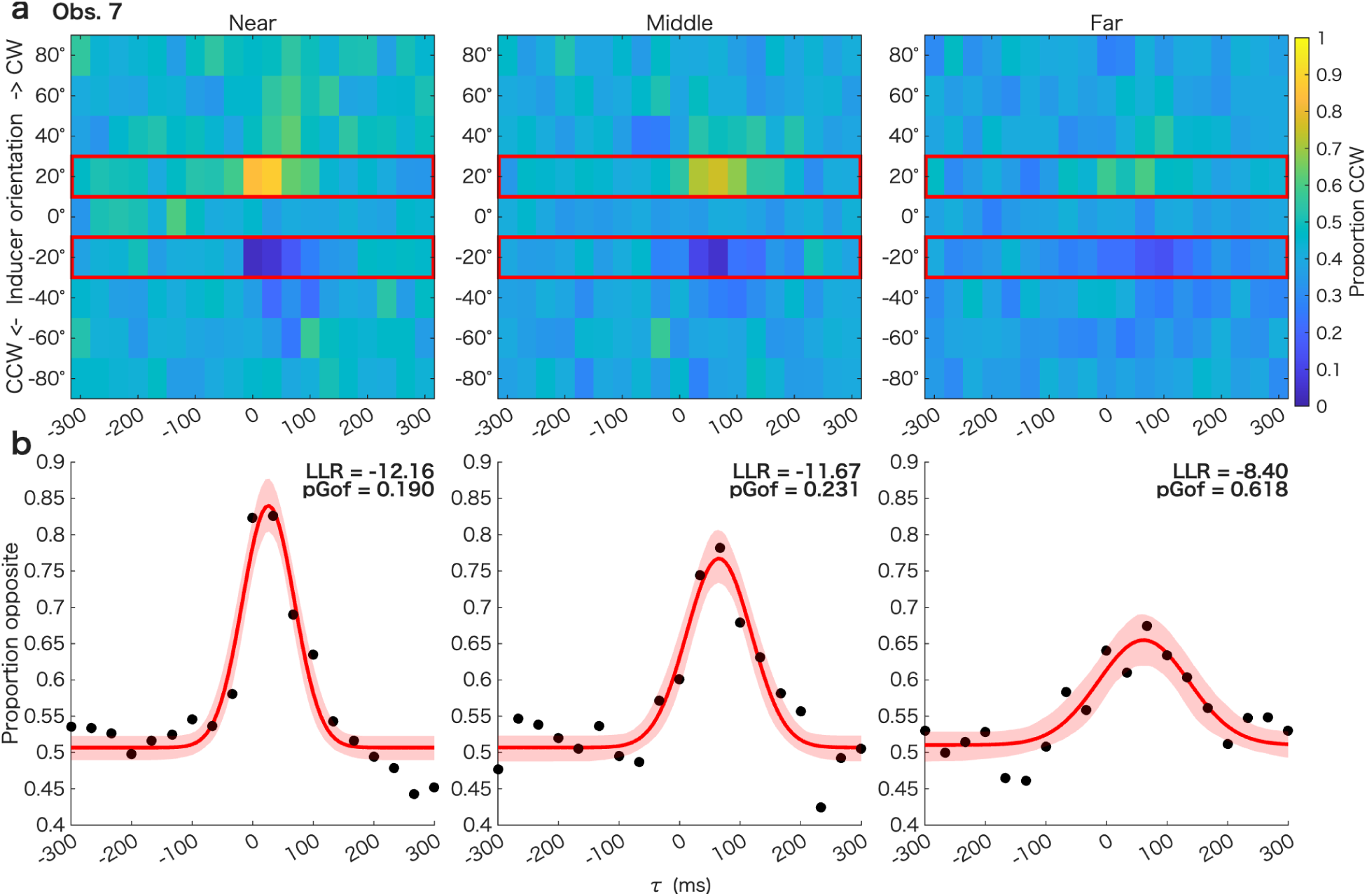
Data for one naïve observer. (a) Correlograms in the near, middle, and far conditions. Each correlogram indicates the proportion of choices by which the observer indicated the horizontal target as appearing counter-clockwise (Proportion CCW) at each combination of inducer orientation (positive = clockwise) and onset time. τ > 0 means inducer onsets after the target onset. (b) Temporal windows of the tilt illusion with ±20° inducers (highlighted with red boxes in panel a) for this observer. Black markers are the actual proportion of trials in which the observer reported the target orientation as tilted against the inducer (“Proportion opposite”), which was derived from the Proportion CW for −20° inducers and the Proportion CCW for +20° inducers. The red curves and shades indicate the best-fit Gaussian functions and their bootstrapped 95% confidence intervals. “LLR” and “pGoF” indicate the log-likelihood ratio of the Gaussian model relative to the saturated model and the p-value for the goodness-of-fit test, respectively. Data for all ten observers are shown in Supplementary Fig. 1. Overall, repulsion peaked when ±20° inducers were presented shortly after the target onset.

As a group-level analysis, we fitted a linear mixed-effects model on Gaussian parameters with observer as a random effect and distance as a fixed effect. Here we report the results when we treated distance as a parametric predictor (2.0, 3.5, and 5.0 dva). The main effect of distance on peak time (β = 8.63 [5.94, 11.32], *t*(27) = 6.58, *p* < .001, *R*^2^ = .358, *BF*_10_ = 3732.5; Fig. 3a), amplitude (β = −0.036 [−0.049, −0.024], *t*(27) = −5.99, *p* < .001, *R*^2^ = .411, *BF*_10_ = 360.7; Fig. 3b), and full width at half maximum (FWHM) (β = 16.82 [10.81, 22.83], *t*(27) = 5.74, *p* < .001, *R*^2^ = .507, *BF*_10_ = 976.8; Fig. 3c) were all significant. In contrast, the main effect of distance on baseline level did not reach significance (β = 0.004 [−0.0004, 0.008], *t*(27) = 1.87, *p* = .073, *R*^2^ = .108, = 2 *BF*_10_ 2.07; Fig. 3d).

**Fig. 3.**
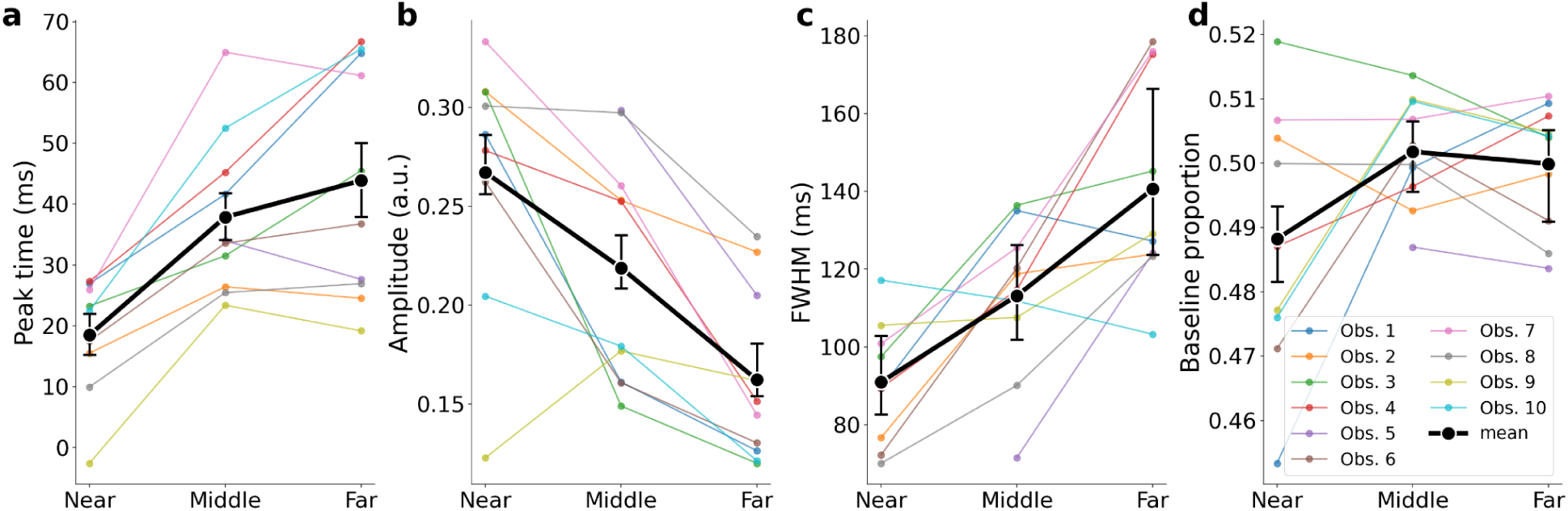
Parameter estimates of the Gaussian temporal window of the tilt illusion when the inducers were tilted ±20° from the horizontal target: peak time. (a), amplitude (b), full width at half maximum (FWHM) (c), and baseline level (d). Coloured lines indicate individual data, and black lines indicate the inter-observer means. Error bars are bootstrapped 95% confidence intervals of the means determined by resampling the proportion opposites per each observer and τ. With increasing target–inducer distance, the temporal window shifted later, became shallower, and broadened.

Pairwise comparisons with Holm’s correction^26^ revealed that the peak time in the near condition was significantly earlier than those in the middle (*t*(17) = 6.43, p < .001, *R*^2^ = .455, *BF*_10_ = 224.7) and far (*t*(17) = 7.17, *p* < .001, *R*^2^ = .463, *BF*_10_ = 312.3) conditions. In contrast, peak time did not significantly differ between the middle and far conditions (*t*(18) = 1.82, *p* = .086, *R*^2^ = .037, *BF*_10_ = 0.94). Importantly, peak time in all three conditions occurred significantly after target onset (near: *t*(8) = 5.66, *p* < .001, *d_z_* = 1.89 [0.75, 2.98], *BF*_10_ = 75.8; middle: *t*(9) = 9.03, *p* < .001, *d_z_* = 2.85 [1.41, 4.28], *BF*_10_ = 2426.5; far: *t*(9) = 7.23, *p* < .001, *d_z_* = 2.29 [1.06, 3.48], *BF*_10_ = 526.1), indicating that an inducer tilted ±20° from the target exerted its strongest repulsive effect when it was presented after the target. Furthermore, the peak time shift across conditions demonstrates that inducers in the farther future contributed to repulsion as the distance increased, but this shift eventually plateaued (Fig. 3a).

Amplitude significantly differed between all the three conditions (near vs. middle: *t*(17) = −2.52, *p* = .022, *R*^2^ = .170, *BF*_10_ = 2.44; near vs. far: *t*(17) = −4.81, *p* < .001, *R*^2^ = .520, *BF*_10_ = 157.3; middle vs. far: *t*(18) = −5.28, *p* < .001, *R*^2^ = .257, *BF*_10_ = 38.6). This amplitude reduction with increasing distance (Fig. 3b) is consistent with previous findings on the spatial tuning of repulsion^27,28^. This relationship could not be fully compensated for by the size scaling of the inducers (see Methods).

FWHM also significantly differed between all the three conditions (near vs. middle: *t*(17) = 3.63, *p* = .004, *R*^2^ = .329, *BF*_10_ = 14.0; near vs. far: *t*(17) = 5.10, *p* < .001, *R*^2^ = .591, *BF*_10_ = 311.2; middle vs. far: *t*(18) = 3.36, *p* = .004, *R*^2^ = .283, *BF*_10_ = 9.60). This widening with increasing distance (Fig. 3c) supports Motoyoshi’s^29^ idea that spatial interactions between orientation detectors involve a greater degree of temporal blur as the detectors become more distant. Intriguingly, the FWHM increased linearly with peak time across conditions and observers (*r* = .537 [.212, .755], *t*(27) = 3.31, *p* = .003, *BF*_10_ = 17.81), consistent with a Weber-like scaling of latency^30^ under the assumption that the FWHM reflects the variance of latency.

To summarize the results, the farther the inducer was from the target, the farther the temporal window of repulsion shifted into the future, becoming flatter and wider.

### Computational modelling favours an on-demand contextual modulation

The distance-dependent shift in the temporal window, by itself, is broadly consistent with the idea that contextual signals require time to propagate through the retinotopic representation. If signals must travel farther to mediate contextual modulation, it would naturally take more time given a constant signal propagation speed. To assess the biological plausibility of the time constant for signal propagation, we estimated the cortical propagation speed from the peak shift data. The speed estimated from the near–middle comparison (∼0.1 m/s) as well as middle–far comparison (∼0.4 m/s) aligns well with the range of unmyelinated horizontal (intracortical) connections within macaque V1^31,32^. This range is also consistent with the mesoscopic cortical travelling waves (0.1–0.6 m/s)^33–35^, which are thought to propagate via horizontal intracortical connections^36^. Together, these agreements in the time constant of physiological signal propagation lend further support to the argument that the tilt illusion is related to neural signals associated with V1^37–39^.

However, the direction of the peak time difference is unexpected if only the distance-dependent delay from the surround to the centre^14,15,23^ is taken into account. The default assumption here is that the surround representation sends out suppressive signals non-specifically to all directions as long as the inducer is present and the process for a central target only *passively* receives outer broadcast signals. Given that the arrival time of a modulatory signal that produces the maximal effect is fixed relative to target onset, a surround whose signal takes longer to arrive at the centre should be presented earlier^17,40,41^. In other words, repulsion should peak earlier for more distant inducers, compensating for the longer propagation delay. And yet, what we found was quite the opposite: more distant inducers exerted their strongest influence when they were presented progressively later relative to the target.

We therefore considered an alternative hypothesis in which contextual modulation operates *on demand*, i.e., initiated by a query signal that propagates from the centre to the surround only when a central target is individuated. A key implication of this account is that repulsion is driven most strongly by inducers presented after the target, because contextual information is made available only after the query signal sent from the centre reaches the surround. Furthermore, under this account, more distant inducers may become increasingly effective when presented later, because their contribution to repulsion should be maximal when their direct response coincides with the arrival of the query signal. As a proof of concept for this on-demand account, we constructed a mechanistic model that extends the dynamic divisive normalization model^25,42^ by incorporating signal propagation delays in centre–surround interactions (Fig. 4).

**Fig. 4.**
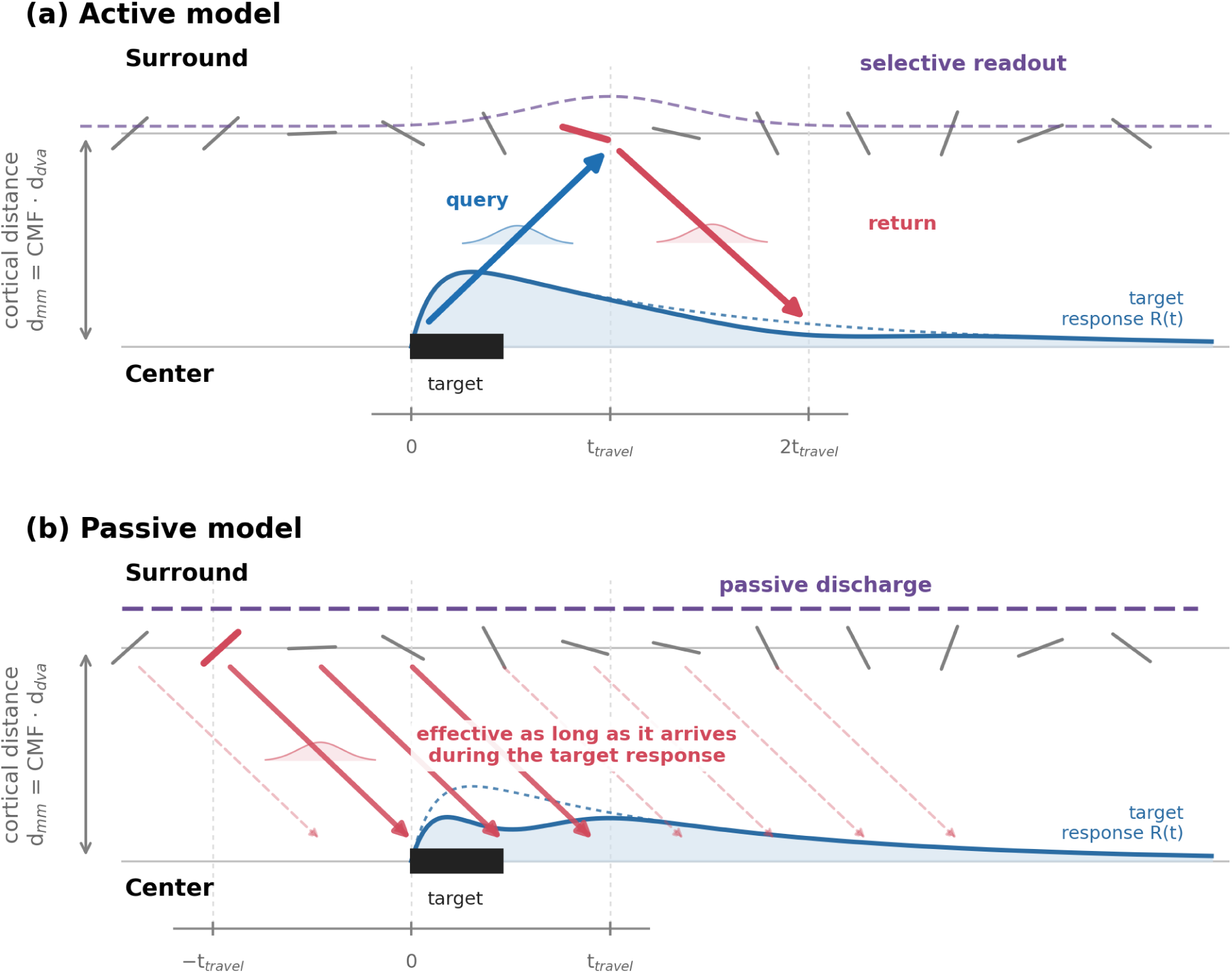
Schematics of the two competing mechanistic models. (a) Active model. A target onset triggers a query on demand that propagates outward for an interval t_travel_ to the surround. A reply signal from the surround in response to the query returns to the target another t_travel_ later and participates in a divisive normalization of the target response function. Because more distant inducers are recruited later, the effective inducers are those presented later in time, causing the temporal window to peak progressively later with increasing distance, as seen in the observed data. (b) Passive model. There is no selective recruitment by a query; every inducer continuously sends its suppression signals inward, reaching the target after a delay t_travel_. A given inducer is effective only if its suppression signal arrives while a target response is present and left immature (solid arrows). Because the suppression must coincide with the target response, the effective inducers are those presented earlier, causing the temporal window to peak progressively earlier with increasing distance, opposite to the observed data. Both panels are space–time diagrams: the horizontal axis is time relative to target onset, and the vertical offset of the two timelines is the cortical distance between the target (centre) and inducer (surround). Bars on the surround timeline denote inducer orientations (refreshed every 33 ms) and the black bar on the centre timeline is the target. The solid blue curve schematizes the target response R(t), the dotted curve the response expected without normalization, and each bell-shaped curve the temporal blur in query/return signals.

More specifically, we formalized and contrasted two model architectures. In the passive model, surround responses continuously propagate toward the target representation with a distance-dependent delay and modulate the target response through divisive normalization whenever the inducer is present (Fig. 4b). In the active model, by contrast, the same contextual signals are able to influence the target representation only when they are recruited by a target-triggered query signal that propagates from the centre to the surround (Fig. 4a).

The active model (Fig. 5a and Supplementary Fig. 2) provided a substantially better fit to the data than the passive model (ΔAIC = 2670; 86.3% vs. 29.0% deviance explained; see Fig. 5b and Supplementary Fig. 3). Importantly, the fitted propagation speed *v*_ℎ_ of the active model fell within a biologically plausible range (0.06–0.16 m/s; mean 0.10 m/s; see Supplementary Table 1), a similar order of magnitude as the conduction velocity of horizontal connections^31,32,34^. When we fit the Gaussian function on the model-predicted correlogram, the qualitative trends of the summary statistics were reproduced as well: the active model reproduced the increase of peak time, the decrease of amplitude and the increase of FWHM, with increasing distance (Fig. 5a). Over the 30 summary statistics (10 observers × 3 distances), the observed and predicted values correlated at r = .88 for peak time, .96 for amplitude and .76 for FWHM.

**Fig. 5.**
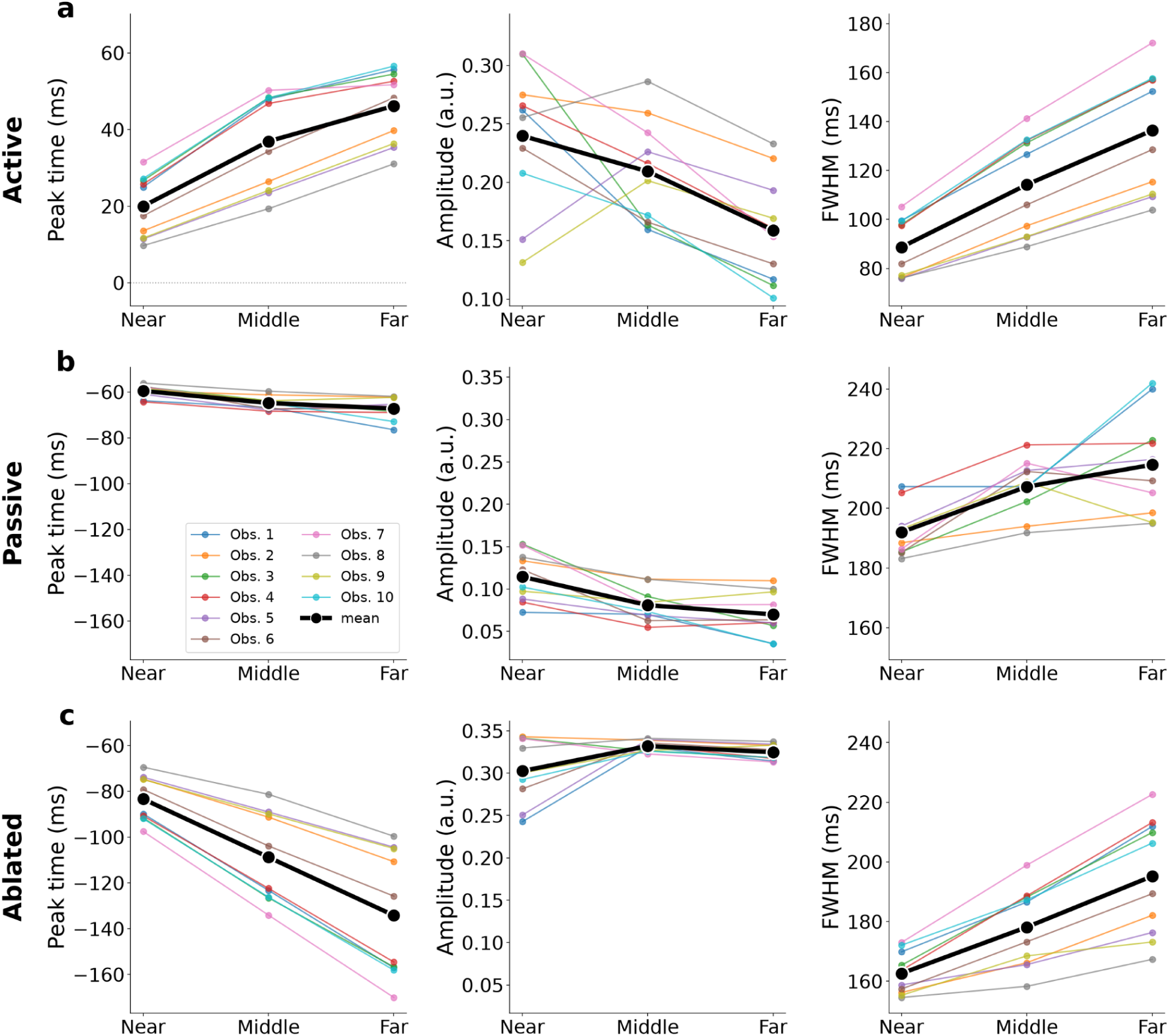
Results of computational modelling. (a) the mechanistic active model, (b) the mechanistic passive model, and (c) the active model with its query components ablated without refitting (ablated model). Columns show the peak time (left; the dotted line marks zero), the amplitude (middle) and the FWHM (right) of a Gaussian fitted to each predicted correlogram, obtained in the same way as for the observed data (cf. Fig. 3). Coloured lines indicate the statistics of the model’s prediction for the individual data, and black lines indicate the inter-observer means. Note that the y-axes in (a) are identical to those of Fig. 3, whereas (b) and (c) share a common y-axis that differs in range and scale from that of (a), for readability. Only the active model reproduced the later peak time with increasing distance that was observed in the data (a; cf. Fig. 3a). The passive model instead predicted a nearly constant, negative peak time (b), and ablating the query from the best-fit active model reversed the peak time shift (c). Therefore, the active model was favoured over the passive one both qualitatively and quantitatively.

Notably, the active model even approached the descriptive Gaussian model in performance (86.3% vs. 89.0% deviance explained; see Fig. 2b and Supplementary Fig. 1), despite the former being constrained by biologically interpretable mechanisms yet using substantially fewer free parameters (K = 53 vs. 120).

By contrast, the passive model (Fig. 5b) not only fit poorly with the data but also failed to reproduce all summary statistics: peak time (r = −.80), amplitude (r = .66) and FWHM (r = .40). Most strikingly, its predicted peak time was nearly constant at about −65 ms (opposite sign to the data) regardless of distance and observer. The constant peak time was achieved by driving the propagation speed *v*_ℎ_ to its upper bound of search range (5.0 m/s; see Supplementary Table 2) for all observers, effectively eliminating the delay of signal propagation between the centre and surround. This boundary solution, which persisted regardless of the search range, reflects a constraint of the passive model architecture: under the assumption of biologically plausible propagation delay, the model inherently predicts *earlier* peak time with increasing distance. Thus, fitting the passive model to the data with *later* peak time with increasing distance forcibly pushes the delay toward zero, but never toward positive values as in the observed data. Consistent with this, removing the query mechanism from the best-fit active model while keeping all remaining parameters fixed (i.e., ablated model) produced an earlier peak time, quite the opposite to the data (Fig. 5c, −344.8% deviance explained). Therefore, the passive model fails exactly because of the absence of the query mechanism.

Taken together, the present findings point to two key temporal properties of the tilt illusion. First, the inducer exerts its strongest influence after target presentation rather than before it. Second, more importantly, the most effective inducer occurs progressively later as the distance increases. Both properties follow from a single query process with a single conduction velocity: inducers become effective only when the query reaches the surround, which places the window after target onset, and the query takes longer to reach a more distant surround, which moves that moment progressively later. The active model therefore provides a parsimonious mechanistic explanation of the observed psychophysical correlograms.

## Discussion

A central computational challenge for visual processing is to provide the best solution as to how information distributed across distant regions of the visual field should be integrated into a coherent percept. Existing models of contextual modulation successfully account for the steady-state influence of surrounding information on perception, but they provide little insight into how these interactions unfold over time. The present psychophysical data and computational modelling suggest that such interactions do not arise through passive propagation of neural activities across the visual field; conversely, contextual information is recruited on demand after a target has been individuated.

Existing models of the tilt illusion are typically formulated as static computations in which the response to a target is normalized by surround responses without explicitly written temporal dynamics. As a consequence, these models define contextual influences only in terms of how the surround affects the target representation, while remaining agnostic about any causal signal flow at the neural circuit level. The poor fit with the model of passive surround-to-centre propagation, together with the successful fit with the active model, suggests that contextual modulation involves signals originating from the target representation in addition to those arriving from the surround.

Such bi-directional interactions have been implemented in some recurrent circuit models to account for contextual phenomena at the neuronal level^15,43,44^. However, linking such dynamic mechanisms to perceptual phenomenology has been challenging because psychophysical measurements typically provide only the final outcome of underlying computations. By combining reverse-correlation analysis with systematic manipulations of spatiotemporal parameters, the present study provided constraints on the directionality of contextual interactions, favouring bi-directional accounts over surround-to-centre (i.e., uni-directional) accounts.

Our proposed active model, assuming bi-directional interactions, is also compatible with physiological dynamics of surround suppression measured in previous studies. These studies, however, essentially fixed the stimulus onset asynchrony between the centre and surround while manipulating their spatial distance^14,15^. Since the modulation delay should grow with distance regardless of whether the interaction is initiated by the centre or surround, such a design cannot distinguish active from passive mechanisms. Thus, the physiological data constrained the neural circuits and timescales of signal propagation and our psychophysics data with the full-range sampling of stimulus onset asynchronies provide an additional constraint on the directionality of these interactions, which has not previously been considered.

The proposed active mechanism presupposes that a target has already been individuated, although not finalized yet, before contextual information is recruited. Therefore, our model may be particularly relevant to contextual processes that modify the visual appearance of an existing object representation, rather than to processes in which contextual interactions themselves contribute to object individuation, such as figure–ground segregation^45,46^ or visual pop-out^47,48^. Such early segmentation processes likely need to operate rapidly in order to guide subsequent visual processing, and may therefore depend more strongly on rapid local interactions or feedback mechanisms than on the long-range sluggish process proposed here^49^.

Iso-orientation surround suppression has been argued to support efficient coding of natural images^12,50–52^. Natural images are highly structured, giving rise to statistical interdependencies among visual features across space. For example, due to the continuity of object contours and surface textures, orientation statistics are typically more similar between nearby locations than between distant locations^53^. Under such conditions, responses to a visual feature can be predicted from its surrounding context, making it advantageous to suppress redundant information, to control the dynamic range, and to emphasize deviations from local expectations. Our findings suggest a further form of coding efficiency beyond the spatial redundancy reduction. Specifically, under the on-demand surround suppression account, computational resources need not be continuously allocated to integrating contextual information across the visual field. Instead, contextual information may be recruited only when the visual system needs to evaluate a target representation in reference to its surroundings.

The implications of the present findings may extend beyond the tilt illusion. Contextual phenomena on visual appearance are observed across a wide range of visual features, as reviewed elsewhere^54,55^. The proposed on-demand recruitment of contextual information may therefore be a more general computational strategy, particularly for contextual effects relying on long-range interactions, such as perceptual filling-in^56,57^, apparent motion^58^, maintenance of object files^59,60^, contour integration^61,62^, perceptual constancy^63^, and perceptual synchrony^64^, where signal propagation and interaction incur substantial temporal and metabolic costs. Consistent with this idea, for example, Cass and Spehar^40^ reported a similar distance-dependent temporal profile for collinear facilitation, a phenomenon in which the detectability of a grating is enhanced by collinear surrounding gratings (i.e., flankers). Specifically, more distant flankers required longer exposure durations before facilitation emerged.

The precise characterization of the temporal window in the present study also provides insights into temporal processing in the visual system. The peak time of approximately 50 ms for the farthest inducer suggests that the conscious perception of repulsion finalizes no earlier than ∼100 ms following target registration. This length of buffer time is consistent with previous studies reporting that extending the inducer duration by up to ±100 ms around the target onset increases the strength of repulsion^65^. The different arrival times of modulatory signals from different distances observed in the present study may underlie the gradual buildup of target orientation representation preceding the final perception of repulsion^66–68^.

In contrast to the relatively coarser temporal resolution at which conscious percepts are considered to be updated^69,70^, our results suggest that the contextual modulation underlying the tilt illusion operates more rapidly. Given that inducer orientations changed every 33 ms, the contextual interactions underlying the repulsion must be capable of sampling surround orientations on at least this timescale, and potentially much faster^22,65^. This distinction of temporal resolution and temporal extent^65^ suggest that temporal precision of computations contributing to contextual modulation exceeds the temporal precision with which those computations become consciously accessible. Such a dissociation is compatible with theoretical accounts proposing that conscious perception is shaped by temporally precise information, even though conscious contents themselves are updated at a coarser temporal resolution^70,71^. Indeed, the temporal evolution of repulsion itself can become consciously perceptible when it unfolds sufficiently slowly. Yuan et al.^72^ demonstrated that a physically static target appeared to swing back and forth when the alternation rate of clockwise and counterclockwise inducers was lower than 6 Hz. Together with the present findings, the visual system may continuously compute contextual influences at a relatively high temporal precision (tens of milliseconds), while conscious perceptual contents are updated more slowly, possibly on the order of hundreds of milliseconds^73,74^.

More generally, the present findings raise the possibility that transient sensory buffers (i.e., visual and informational persistence) play a computational role beyond mere retainment of information after stimulus offset. Long-range contextual interactions inevitably require time because information must propagate between distant retinotopic representations. Maintaining sensory representations in an unfinalized state may therefore allow the visual system to postpone conscious perception until these interactions have been completed. From this perspective, apparently *postdictive* relationships^75–81^, in which a future event apparently influences the contents of a preceding event, may be viewed as a consequence of such a strategy that enables integration of spatiotemporally extended information.

At the same time, the rapid changes in inducer orientation in the present paradigm constrain the class of contextual processes directly characterized by our measurements. Some contextual effects are known to operate on timescales substantially slower than the 33-ms inducer updates used here. For example, the rod-and-frame effect (i.e., a vertical rod appearing tilted when placed inside a tilted frame) is stronger at relatively low temporal frequencies^82,83^, with some evidence suggesting that contextual influences emerge at relatively late, possibly post-perceptual stages of processing^84^ and involve the integration of visual and vestibular information in constructing a global reference frame for orientation^85,86^. These properties suggest that the mechanisms underlying the rod-and-frame effect are unlikely to be identical to the rapid, relatively local contextual interactions examined here.

Nevertheless, its temporal dynamics is still compatible with the principle of active contextual processing. Peterson and Dassonville^41^, for example, found that a tilted frame could influence the perceived orientation of a briefly presented rod even when the rod preceded frame onset by approximately 180 ms. They attributed this apparent postdictive effect to differential processing latencies, whereby orientation judgment of the rod remains ongoing when the subsequent frame begins to alter its reference frame. An active account goes beyond this description of the time course by specifying the flow of information underlying contextual processing: target registration initiates contextual recruitment, whose outcome is subsequently integrated into the target judgment, although whether the neural timescale of such information exchange can account for the slower dynamics of the rod-and-frame effect needs to be tested. In this way, despite differences in spatiotemporal scale, active contextual recruitment has potential to represent a general computational principle across distinct forms of contextual modulation.

To our knowledge, this is the first study to develop a dynamic mechanistic model of the tilt illusion, wherein signal propagation delays between the centre and surround are incorporated explicitly. Despite its biological plausibility, our mechanistic model is intentionally simplified and does not include all components that may contribute to or interact with the tilt illusion, as doing so would introduce substantial parameter indeterminacy. First, surround suppression may partly arise from feedback signals originating in extrastriate visual areas^7,87,88^, which was not explicitly modelled. Bair et al.^14^, by which our study was inspired, found that suppression from more distant surrounds tended to arrive later, but also the inferred propagation speed (∼1 m/s) was faster than expected from long-range horizontal connections alone. They instead proposed that rapid suppression from distant surrounds could involve feedback from extrastriate neurons with large receptive fields. Although our results do not rule out a contribution of feedback signals to the tilt illusion, such a contribution cannot by itself account for the systematic, distance-dependent shift in peak times. The presence of feedback therefore does not alter our main conclusion that the observed distance-dependent delay points to slow horizontal propagation.

Second, our observers may have suffered from the phenomenon of crowding (i.e., a central visual object becoming harder to discern when flanked by similar objects), particularly in the near condition, because the inducer was presented within the known critical spacing for crowding^89–91^. Crowding is also known to exhibit temporal tuning centred around target onset^92^. Under crowding the target orientation is not simply lost but could be replaced by, or averaged with, the orientation of flanking stimuli^93,94^. On the trials where this happens, the observer effectively reports the inducer orientation, which counts as a choice *toward* the inducer (i.e., an attractive bias) and therefore pushes the proportion opposite below 50%. However, because crowding is expected to be strongest near target onset, any such attractive component would artifactually shift the measured peak in the near condition away from zero. Such a displacement would move the near-condition peak towards those of the middle and far conditions, thereby reducing the distance dependence we report, rather than creating it. Therefore, crowding cannot account for the observed pattern of results (see also Supplementary Notes and Supplementary Fig. 4).

Third, because the width of the annular inducer increased with target–inducer distance, the number of neuronal units activated by the inducer and potentially contributing to surround suppression may have differed across conditions. However, the effective strength of contextual modulation likely reflects a combination of several factors, including the number of recruited neural units, the spatial extent of attention, and the intrinsic strength of surround suppression. Our model assumes that these factors act as a gain on the contextual effect, scaling the amplitude of the correlogram while leaving the temporal parameters unchanged. Each distance was therefore given its own amplitude scaling factor that absorbs these contributions altogether, while the peak times on which our conclusions rest are unaffected by this scaling.

In summary, the present study used a psychophysical reverse correlation technique to demonstrate that the temporal window of a repulsive tilt illusion depends on the target–inducer distance. The observed peak time differences across different distances are consistent with the neural latency of iso-orientation surround suppression in the primary visual cortex. The direction of the peak time shift suggests a temporally unfolding mechanism that is initiated only after the target is registered. On-demand recruitment of spatiotemporally extended contextual information may reflect a general strategy by which the visual system allocates costly long-range computations efficiently in time, with implications for a broad class of contextual phenomena that shape perceptual appearances.

## Methods

### Observers

Nine adults who were naïve to the purpose of the study and the first author (six men and four women; aged 21–28 yrs) having normal or corrected-to-normal vision participated. All had prior experiences with psychophysical experiments. The study was approved by the ethics committee of the Graduate School of Humanities and Sociology at the University of Tokyo and conducted with adherence to the Declaration of Helsinki. We obtained a written informed consent from each observer prior to the experiment.

### Apparatus

The experimental programs, written in MATLAB R2019b (MathWorks, Natick, MA, USA) and Psychophysics Toolbox Version 3^95,96^, were executed on a computer (Dell Precision with Ubuntu OS ver. 18.04.5). Visual stimuli were displayed on a gamma-corrected 22-in. CRT monitor (Mitsubishi Electric RDF223H; 1600× 1200 pixels; 60 Hz) via a graphics board (AMD Radeon Pro WX 5100). Black cardboard occluded the monitor edge to prevent it from serving as a reference frame. The observers viewed the stimuli binocularly, resting their heads on a chin-and-head rest positioned 57cm from the monitor. Throughout the experimental sessions, the gaze of one eye was recorded using an eye tracker (EyeLink 1000, SR Research) with a sampling rate of 250 Hz.

### Stimuli

Stimuli were presented on a grey background (31 cd/m^2^). A black bullseye (0.8 dva × 0.8 dva) was used as a fixation point on which the observers were instructed to maintain fixation throughout each session, except during the probe and inter-trial intervals. As the inducer, an annulus-shaped sine-wave grating was presented 8.0 dva below the fixation point, a peripheral location at which repulsion is stronger than in central vision^97^. The average radius of the grating *d_dva_* was 2.0, 3.5, and 5.0 dva in the near, middle, and far conditions, respectively. The contrast of the annulus was 0.99 but tapered at the edges:

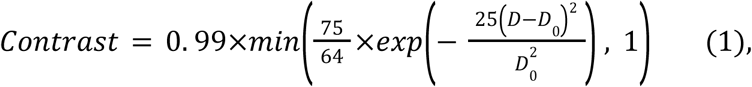

where *D* indicates the distance from the centre and *D*_0_ indicates the average radius. Accordingly, annulus width was proportional to *D*_0_. This size scaling was intended to compensate for the decrease in repulsion with increasing target–inducer distance^27,28^. We used Gabor patches (contrast: 0.99; SD of Gaussian envelope: 0.31 dva) for both the target and probes described below. The spatial frequencies of the inducer, target, and probe were 2 cpd, with their phases randomly chosen in each trial.

### Procedure

The near, middle, and far conditions were intermingled in a random order from trial to trial. Each trial began with the onset of the fixation point. After fixation was maintained for 800 ms, nineteen inducers were sequentially presented. Each inducer had a random orientation, chosen from 0°, ±20°, ±40°, ±60°, ±80° relative to the horizontal. Throughout, positive angles denote clockwise tilts from horizontal. Each orientation lasted for 33 ms with no inter-stimulus interval. Alongside the tenth inducer, a horizontal target was flashed for 33 ms at the centre of the annulus. By fixing the target onset timing this way, we maximized the target’s temporal predictability, thereby minimizing the differential latency between target and inducer^18,98,99^.

The sequence of inducers was followed by a 300-ms blank period and a subsequent probe period, during which two probes, a randomly chosen one of them tilted +5° and the other −5°, were presented 10 dva to the left and right of the target location. The observers were asked to indicate which probe orientation better resembled the target orientation, with no feedback provided. Each choice was followed by a vacant screen, and the next trial began after 1000 ms (Fig. 1).

To maintain engagement, the target was physically tilted either +10° or −10° in catch trials (21%) that appeared in a random order within each session.

In each session, the observers conducted 57 trials (including 12 catch trials) for each of the near, middle, and far conditions. Trial order was fully randomized, with a short break after every 57 trials. Within each session, the inducer of each orientation equally frequently occurred for both the main and catch trials. Eight observers completed 33 such sessions, such that the number of trials associated with each cell in the correlogram for each observer became 165 (Fig. 2), which was 10% larger than that in Mareschal and Clifford’s study^22^, to compensate for the expected loss of some trials due to fixation errors. Due to calibration failures of eye-tracking, the two remaining observers completed 20 and 24 sessions, respectively.

### Data analysis

First, trials (5.3%) in which gaze positions deviated by > 2.0 dva during the inducer sequence were excluded from further analyses.

Based on the inducer sequence and the choice in each trial, we constructed a correlogram between them for each condition and each observer (Fig. 2). The procedure followed the one used by Mareschal and Clifford^22^. In the correlogram, each column indicates the stimulus onset asynchrony (τ) between the target and inducer, while each row indicates inducer orientation. Each cell indicates the proportion of trials in which the (physically horizontal) target was judged as tilted CCW. Following Mareschal and Clifford’s study^22^, we focused on ±20°, the optimal orientations for inducing repulsion. By flipping the proportion CCW for −20° inducers and merging it with the proportion CCW for +20° inducers, we obtained the “proportion opposite,” namely the proportion of trials in which the target was judged as tilted against the inducer orientation^66^. If a given inducer had no influence, a proportion opposite of 0.5 was expected by chance. A higher value indicated a stronger contribution of the inducer at that τ to repulsion.

The proportion opposite as a function of τ was fitted with a Gaussian function, separately for each condition and each observer:

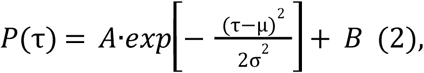

from which the full width at half maximum (FWHM) was derived:

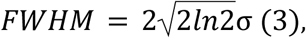

where σ, µ, *A*, and *B* were all free in the four-parameter model.

Model parameters were estimated by the maximum likelihood method, and the model fit was assessed with a likelihood-ratio test^100^. One observer’s data in the near condition failed to reach the goodness-of-fit criterion (*p* = .010, likelihood ratio test) and was therefore excluded from further analyses, although including these data did not alter the statistical conclusions.

### Estimation of signal propagation speed

To estimate the propagation speed underlying the peak shift of the temporal window, we first converted the target–inducer distance into cortical millimeters using an approximation of the cortical magnification factor (CMF) of macaque V1^5,14^:

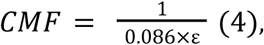

where ε is the eccentricity in dva. For a target at 8 dva, CMF ≈ 1.45 mm/dva, yielding a cortical distance of 2.18 mm for a 1.5 dva separation between two adjacent inducers. The propagation speed *v* was then estimated as:

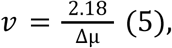

where Δµ is the mean difference in the peak time between conditions (near–middle or middle–far).

Note that we applied the CMF at the target eccentricity regardless of inducer eccentricity, adopting a mid-range estimate from a physiological perspective. We simplified it in this way deliberately because the estimate of CMF has heterogeneity across studies even at the fixed eccentricity. At 8 dva, linear CMF values for V1 span from ≈0.95 mm/dva (macaque, power-law fit of Van Essen et al.^101^) to ≈1.74 mm/dva (human fMRI, Duncan & Boynton^102^), bracketing our value within less than a factor of two. Because *v* scales linearly with the CMF, adopting any of these alternative estimates would change *v* by at most ∼2-fold and would therefore leave the estimated speed within the range reported for horizontal (intracortical) connections in V1.

### Computational modelling

#### Model overview

We formalized mechanistic models of biologically plausible interactions that might be considered the underpinnings of contextual modulation that we observed in the form of correlograms. The active model comprises a cascade of four neural processing stages: (i) population coding of orientations among orientation-selective channels at both the target and inducer locations, (ii) dynamic divisive normalization of the population response to the target by an on-demand recruitment from the surround stream, with centre–surround communication delays scaling with cortical distance, (iii) a population-vector decoder that converts an instantaneous normalized population response into an evidence signal, (iv) a linear integrator that accumulates evidence signals over time and maps the total signed evidence onto a choice probability. All dynamics are first-order ordinary differential equations integrated with Euler steps of 1 ms. We set 5 free parameters for each of our 10 observers: three gain parameters (α*_near_*, α*_mid_*, α*_far_*) governing the suppression strength at each distance, a horizontal conduction speed (*v_ℎ_*) that determines the target-to-inducer travel time, and a lapse parameter (π). We set 3 additional free parameters constrained across observers: a scaling parameter associated with orientation discrimination threshold (σ*_M_*) in psychophysical performance, a coefficient of variation (*k*) that allows travel time to obey a Gaussian function of a certain dispersion, and a latency (*t_reg_*) between target onset and the departure of the query, yielding 53 free parameters (5 × 10 + 3) in total. Other parameters were fixed to values reported in the previous literature or fixed for convenience.

#### Orientation selectivities

At both the target and inducer locations, orientation is represented by a set of 12 channels with preferred orientations θ spaced uniformly over (−90°, 90°]. Orientation tuning is modelled as a Gaussian function of the circular difference of preferred θ*i* and stimulus orientations Θ.

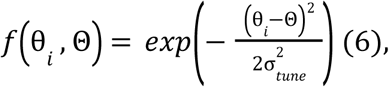

where orientation tuning width is σ*_tune_* = 20° (see ref.^12^)

#### Direct responses to the target and inducer

The excitatory drive *E_c_*_,*i*_(*t*) for channel *i* to the target rises during target presentation (33 ms) and decays thereafter, with a single time constant τ*_E_* = 50 ms. The exponent is set at *n* = 1. 5 following Chapman and Denison^42^.

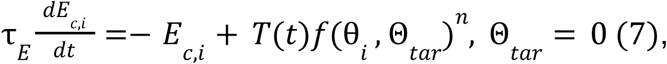

where *T*(*t*) = 1 if *t* ∈ [0, 33), and 0 otherwise.

The excitatory drive *E_s_*_,*i*_ (*t*) for channel *i* to the inducer takes the same form^42^.

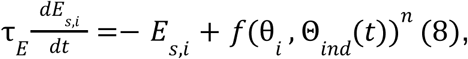

where Θ*_ind_* (*t*) indicates the orientation of the inducer in the currently active time slot at the surround.

#### Query-triggered communication window

We modelled contextual modulation as an *on-demand* communication between the centre and surround. A target onset initiates a query that propagates to the surround; on arrival, the query gets a reply out of the surround, which then returns to the centre and participates in divisive normalization of the target response (Equation 15 below). The query does not reach the surround at a single instant: variability in conduction time disperses its arrival over a finite window, thereby providing the neural origin of the temporal blur normally observed in human psychophysical data such as our correlograms. The query’s arrival window is described as a Gaussian function:

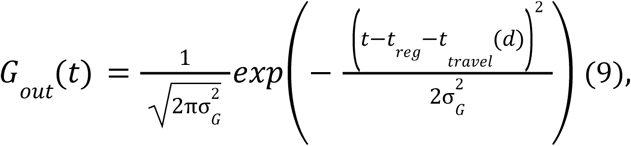

where *G_out_*(*t*) has a unit area and peaks at the arrival of query *t* = *t_reg_* + *t_travel_*(*d*); σ*_G_* controls the width of the window:

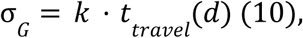

where the one-way conduction delay is determined by the cortical distance *d_mm_* divided by horizontal propagation speed *v*_ℎ_.

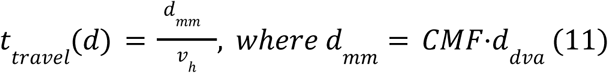

The latency *t_reg_* is the interval between the target onset and the departure of the query, and it represents the time needed for the target to be registered as an event worth querying.

#### Surround recruitment

Each query’s arrival at the surround location evokes a reply signal, that is, the surround response at that time, and thus the distribution of reply signals is described as the pointwise product between the query distribution and the surround stream:

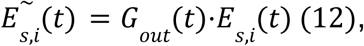

The reply signals then propagate back to the target’s normalization pool, taking a further conduction time *t_travel_*(*d*). Because this return propagation is also temporally dispersed, it is modelled as a convolution with a kernel *G_in_* (*t*) of unit area that has the same Gaussian form and the same width as *G_out_*(*t*), but that peaks at a further conduction time *t_travel_*(*d*) after the query has arrived at the surround:

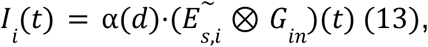

such that the query and the reply together span a round trip, with its expected time of return being 2 · *t_travel_* (*d*). The gain α(*d*) controls the strength of surround suppression. We allowed it to vary at each of the three distances *d*, as the distance dependence of tilt illusion size was out of our experimental control; for example, attentional focus might have differed for different distances and for different observers.

#### Dynamic divisive normalization

The suppressive drive pools the excitatory drive of the target and the reply signal from the surround, and is passed through another temporal filter such that suppression lags excitation. This lag allows for divisive normalization to reproduce typical biphasic temporal receptive fields of sensory neurons^42^.

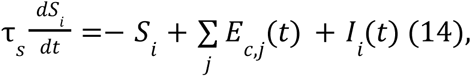

where the suppressive time constant τ*_s_* is set equal to τ*_E_* (50 ms). The divisively normalized response *R_i_*(*t*) for channel *i* to the target is given by:

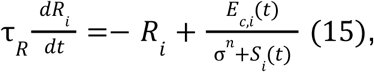

where τ*_R_* = 52 ms and σ = 0. 1, following parameter choices in previous studies^25,42^.

#### Accumulation of evidence from population vectors

To obtain momentary evidence for CW versus CCW choices, we quantified the population vector across orientation-selective channels^103^. However, rather than using the population vector itself to estimate the *perceived* orientation, we extracted only its second-harmonic sine component corresponding to the CW–CCW decision axis *m*(*t*) and linearly accumulated it over time. This formulation preserves the instantaneous magnitude of sensory evidence while avoiding disproportionate influence from time points at which overall population activity has largely decayed toward baseline but still retains residual directional bias.

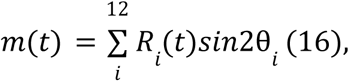

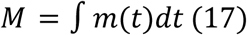

This evidence accumulation is tentatively truncated at the end of the inducer sequence (333 ms after target onset), which our psychophysical task cannot constrain, because overt choices were postponed until the probes appeared. However, we verified that the conclusion of the model comparison did not change even if accumulation extended until *m*(*t*) had decayed to zero.

#### Choice proportions

The accumulated evidence *M* is finally mapped onto a choice probability via a cumulative Gaussian Φ(·) with a scale parameter σ*_M_* associated with orientation discrimination threshold.

For the middle and far conditions,

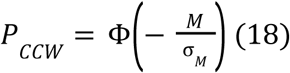

In the near condition, a few observers showed a baseline shift of proportion opposite, which we attributed to a crowding-induced decision strategy whereby, on a fraction π of trials, the observer who missed the target orientation chose between CW and CCW not on a random basis but on the basis of the statistics of inducer orientations. For a simple implementation, in such cases our model compares the count of +20° inducers (*n_CW_*) and the count of −20° inducers (*n_CCW_*) in the sequence and reports the orientation polarity of the most frequent ones:

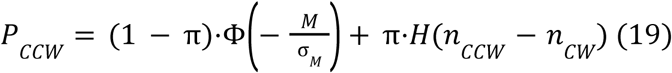

where H(⋅) is the Heaviside step function. This explains why the baseline shifts seen in a few observers unexpectedly persisted in the far past and in the far future relative to the target onset (see also Supplementary Notes).

#### Model optimizations

Model parameters were fit to the choice data by the maximum-likelihood estimation. The objective function was the binomial log-likelihood aggregated over all timeslots, for the +20° and −20° inducer conditions at all distances:

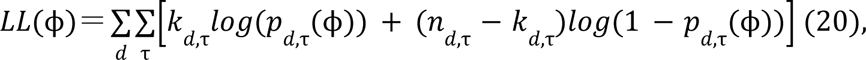

where *k_d_*_,τ_ is the number of choices against the inducer, *n_d_*_,τ_ is the total number of trials, and *p _d_*_,τ_(**φ**) is the mean of the model-predicted “proportion opposite” across trials.

Free parameters were bounded as follows: α ∈ [10^−2^, 10^4^], *v*_ℎ_ ∈ [0. 02, 5. 0] m/s, σ*_M_*∈ [0. 01, 10. 0], π ∈ [0, 1], *k* ∈ [0. 1, 10], *t_reg_*∈ [0, 150] ms.

#### Model comparisons

We compared the fit of the following three models: (i) The descriptive model fitted Gaussian directly on the obtained temporal window without any mechanistic components (the number of free parameters K = 120; see *Data analysis* section); (ii) The mechanistic active model described above (K = 53); (iii) The mechanistic passive model, which was identical to the active model except that surround information was assumed to be continuously available rather than recruited by a target-triggered query. This assumption was implemented by fitting the same model to the data except that *G_out_* (*t*) ≡ 1. The jittered “return” *G_in_* was retained in the passive model, such that the active and the passive models differ only in whether the surround was read out by a target-triggered query. The passive model was fitted with one fewer free parameters (K = 52) because *t_reg_*, a query property, was undefined. In addition, to confirm that the passive model’s poor fit stemmed from the absence of query dependence of the centre-surround communication rather than from its poorly fitted parameter values, we examined (iv) the ablated model by setting *G_out_*(*t*) ≡ 1 in the active model, while all remaining parameters were held fixed without refitting.

Goodness of fit was summarized by the total log-likelihood (LL) and the Akaike information criterion:

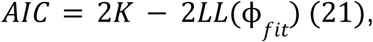

where **φ***_fit_* is the best-fit set of parameters. The formal model comparison between the active and passive models assessed ΔAIC between models, with ΔAIC > 10 taken as decisive evidence for the better-fitting model^104^. To quantify how well each model accounts for the data on an absolute scale, we additionally report the proportion of deviance explained,

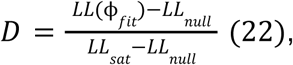

where *LL_null_* is a chance model (0.5 proportion for every cell) and *LL_sat_* is a saturated model in which each cell takes its empirical proportion, i.e., the non-parametric upper bound on fit.

## Acknowledgments

This work was supported by JSPS KAKENHI 22KJ0555, 24KJ0233, 25K18957, 25K00896, 18H05523, 23H01052, and 23K25749. We thank Dr. Hakwan Lau, Dr. Qi Lin and Dr. Ali Moharramipour for insightful comments on earlier drafts of the manuscript.

## Author contributions

Conceptualization: T.N. Data curation: T.N. Formal analysis: T.N. Funding acquisition: T.N., L.O., K.M., and I.M. Investigation: T.N. Methodology: T.N. and I.M. Project administration: T.N. and I.M. Resources: I.M. Software: T.N. Supervision: I.M. Validation: T.N. Visualization: T.N. Writing – original draft preparation: T.N. and I.M. Writing – review and editing: T.N., L.O., K.M., and I.M.

## Data availability

The raw psychophysical data obtained in this study are available in the GitHub, https://github.com/Tomnakam/on-demand-surround-suppression. The raw eye-tracking data is available at Zenodo, https://doi.org/10.5281/zenodo.20780576

## Code availability

All code for the analyses and computational modelling is available in the GitHub, https://github.com/Tomnakam/on-demand-surround-suppression.

