## Supplementary Information for "On-demand surround suppression explains the spatiotemporal window of the tilt illusion"

### Supplementary Notes

In the main text, we report the four-parameter descriptive model (amplitude, peak time, FWHM, and baseline). Here we report a reduced three-parameter model in which the baseline was fixed at 50% (chance). The three-parameter model reproduced the same distance effects (increasing peak time, decreasing amplitude, increasing FWHM; all $p < .001$), and thus our conclusions do not depend on the choice of models. Fixing the baseline parameter actually resulted in poorer fits (i.e., higher AIC values) for the majority of observers, particularly in the near condition (Supplementary Fig. 4). The 95% bootstrapped confidence intervals confirmed that four observers exhibited slightly (< 5%) but significantly lower baseline levels than 50%, whereas one observer exhibited a significantly higher value (51.9%).

This unexpected lower-than-chance baseline level in the near condition is unlikely to reflect perceptual orientation attraction[1], which typically occurs when the orientation difference between the target and inducer is large (e.g., 75–80°). Instead, we suspect that the observers' decision might have been biased toward the inducer orientation when the target orientation became uncertain, possibly due to crowding (see Methods). Although the exact decision strategy cannot be determined solely from our data, we tentatively simulated the observers' choices under the assumption that the observers focused on the orientations near the discrimination boundary (i.e., –20°, 0°, and +20°) and simply chose toward the inducer orientation (+20° or –20°) that was presented more frequently in a given trial. We then derived a correlogram for each observer by simulating the exact inducer presentation sequence from their empirical experiment (excluding catch trials and trials with fixation errors, as in the main analyses) and calculated the proportion opposite for ±20° inducers. The resulting proportions ranged from 25.0% to 36.4% (mean = 30.5%) in the near condition and no systematic relationship with $\tau$ was observed, consistent with the empirical data. Therefore, relying on this decision strategy in some proportion of trials could account for the apparently attractive bias. Motivated by this, we incorporated the same strategy into our mechanistic model as a near-condition-specific lapse parameter ($\pi$), which turned out to improve the fit ($\Delta$AIC = 120; see also Methods).

### Supplementary Figures

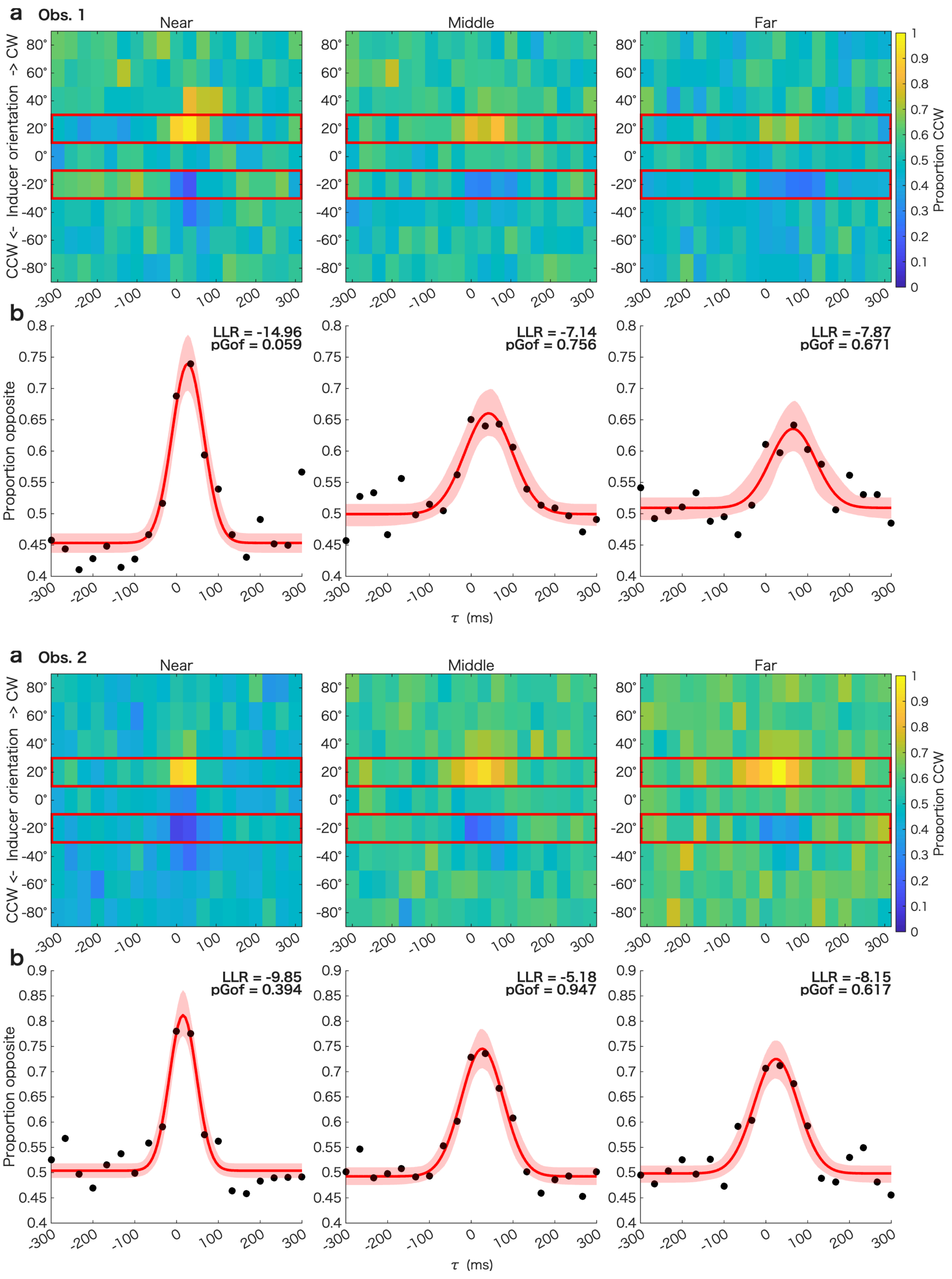

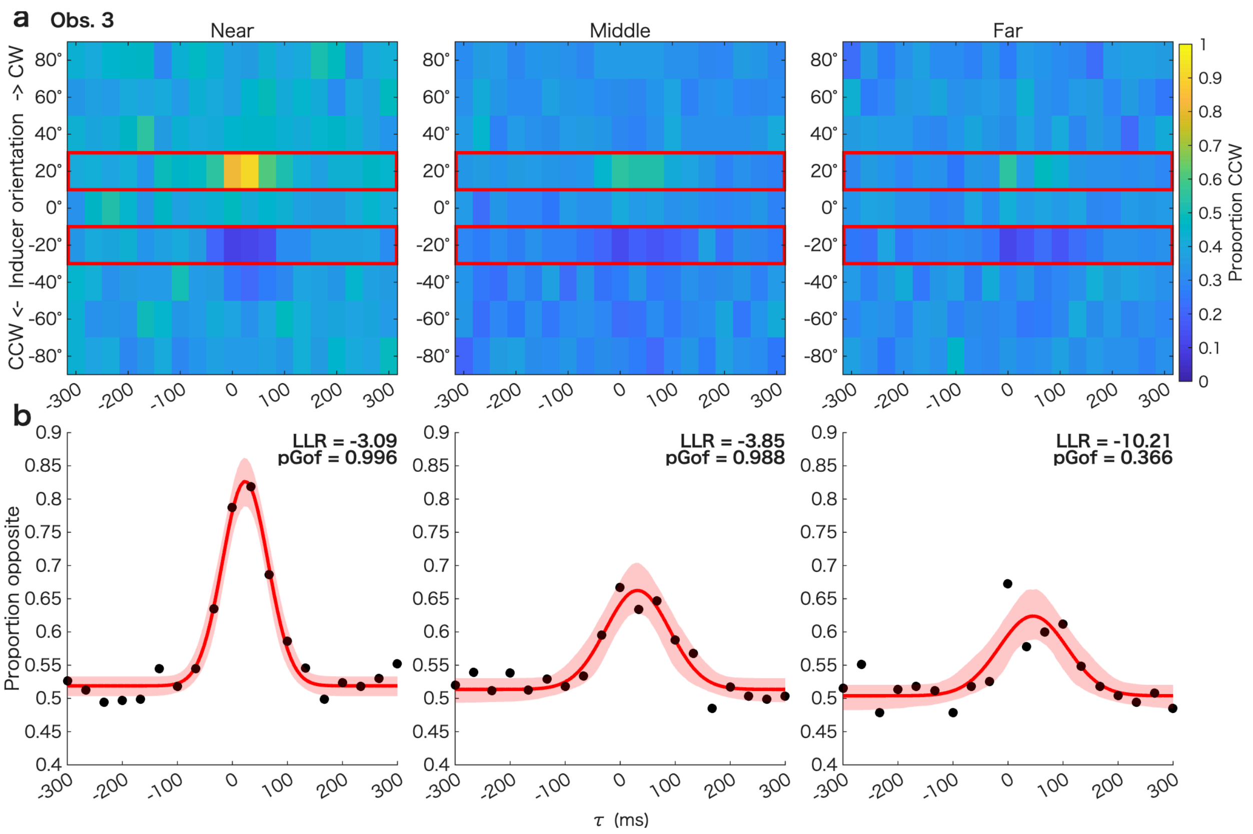

a Obs. 3
Near
Middle
Far
CCW <- Inducer orientation -> CW
80°
60°
40°
20°
0°
-20°
-40°
-60°
-80°
-300 -200 -100 0 100 200 300
Proportion CCW
0 0.1 0.2 0.3 0.4 0.5 0.6 0.7 0.8 0.9 1
b
Proportion opposite
0.4 0.45 0.5 0.55 0.6 0.65 0.7 0.75 0.8 0.85 0.9
LLR = -3.09
pGof = 0.996
LLR = -3.85
pGof = 0.988
LLR = -10.21
pGof = 0.366
τ (ms)


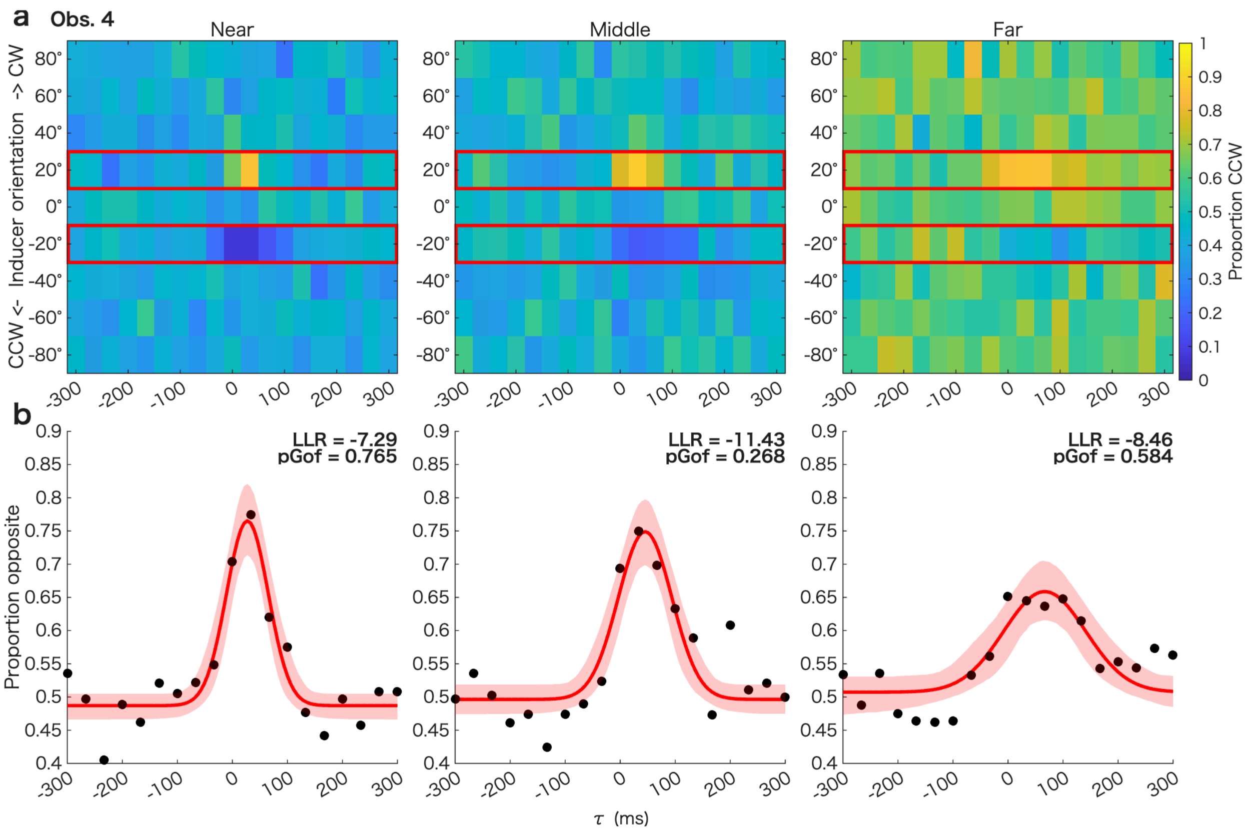

a Obs. 4
Near
Middle
Far
CCW <- Inducer orientation -> CW
80°
60°
40°
20°
0°
-20°
-40°
-60°
-80°
-300 -200 -100 0 100 200 300
Proportion CCW
0 0.1 0.2 0.3 0.4 0.5 0.6 0.7 0.8 0.9 1
b
Proportion opposite
0.4 0.45 0.5 0.55 0.6 0.65 0.7 0.75 0.8 0.85 0.9
LLR = -7.29
pGof = 0.765
LLR = -11.43
pGof = 0.268
LLR = -8.46
pGof = 0.584
τ (ms)

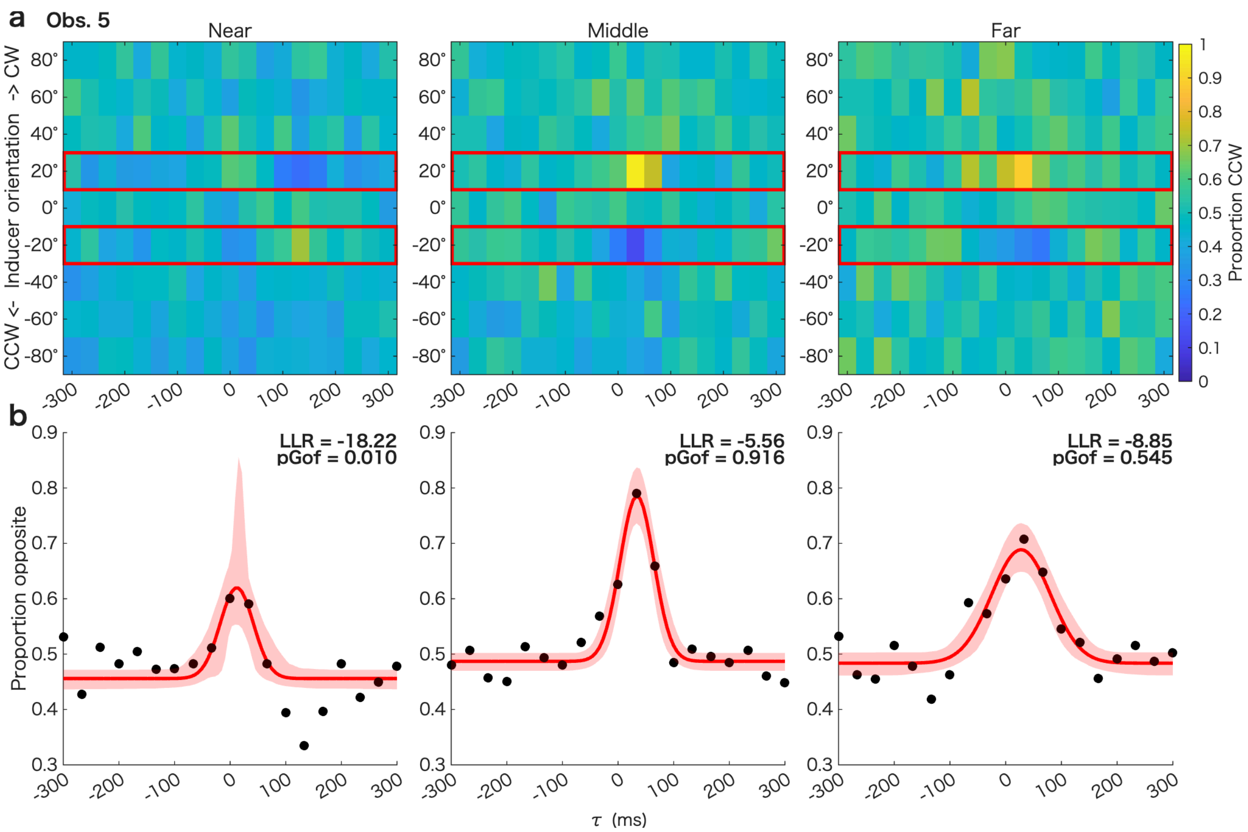

a
Obs. 5
Near
Middle
Far
CCW <- Inducer orientation -> CW
Proportion CCW
b
LLR = -18.22
pGof = 0.010
LLR = -5.56
pGof = 0.916
LLR = -8.85
pGof = 0.545
Proportion opposite
τ (ms)


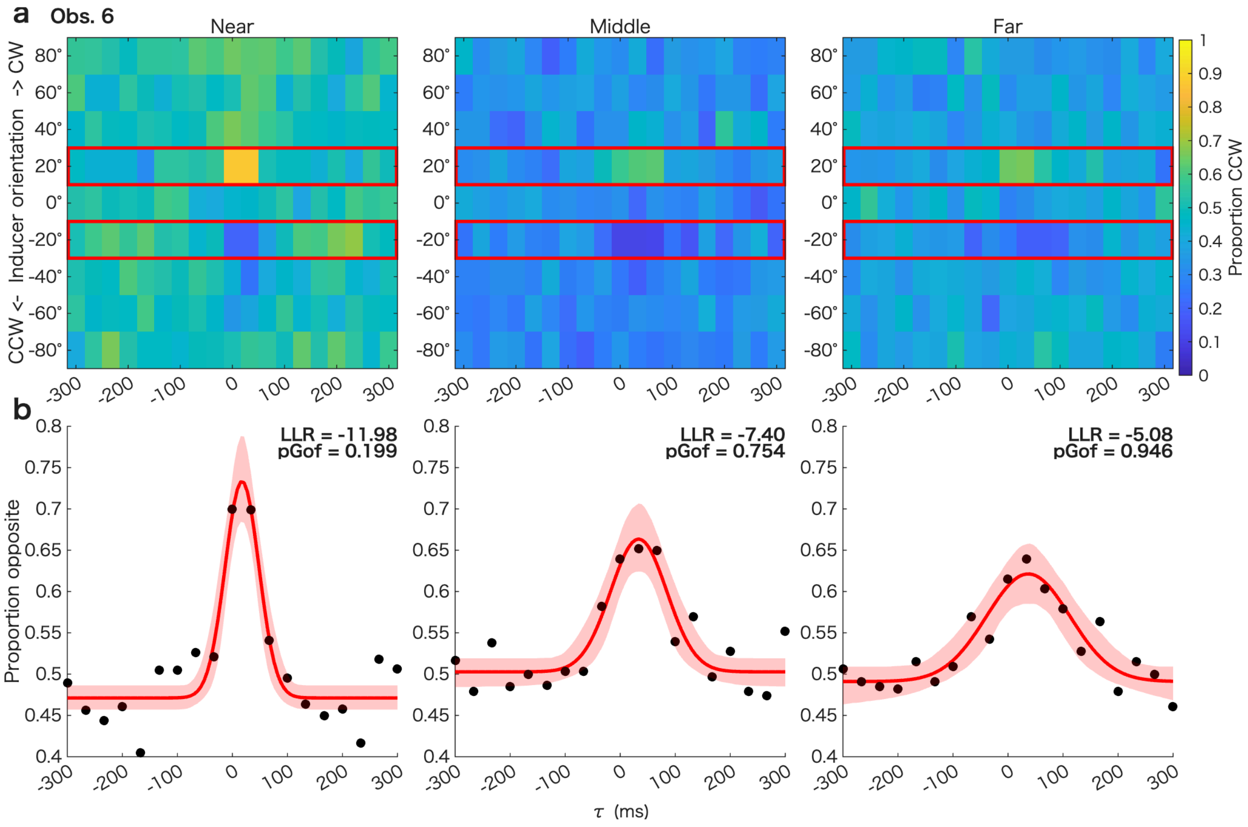

a
Obs. 6
Near
Middle
Far
CCW <- Inducer orientation -> CW
Proportion CCW
b
LLR = -11.98
pGof = 0.199
LLR = -7.40
pGof = 0.754
LLR = -5.08
pGof = 0.946
Proportion opposite
τ (ms)

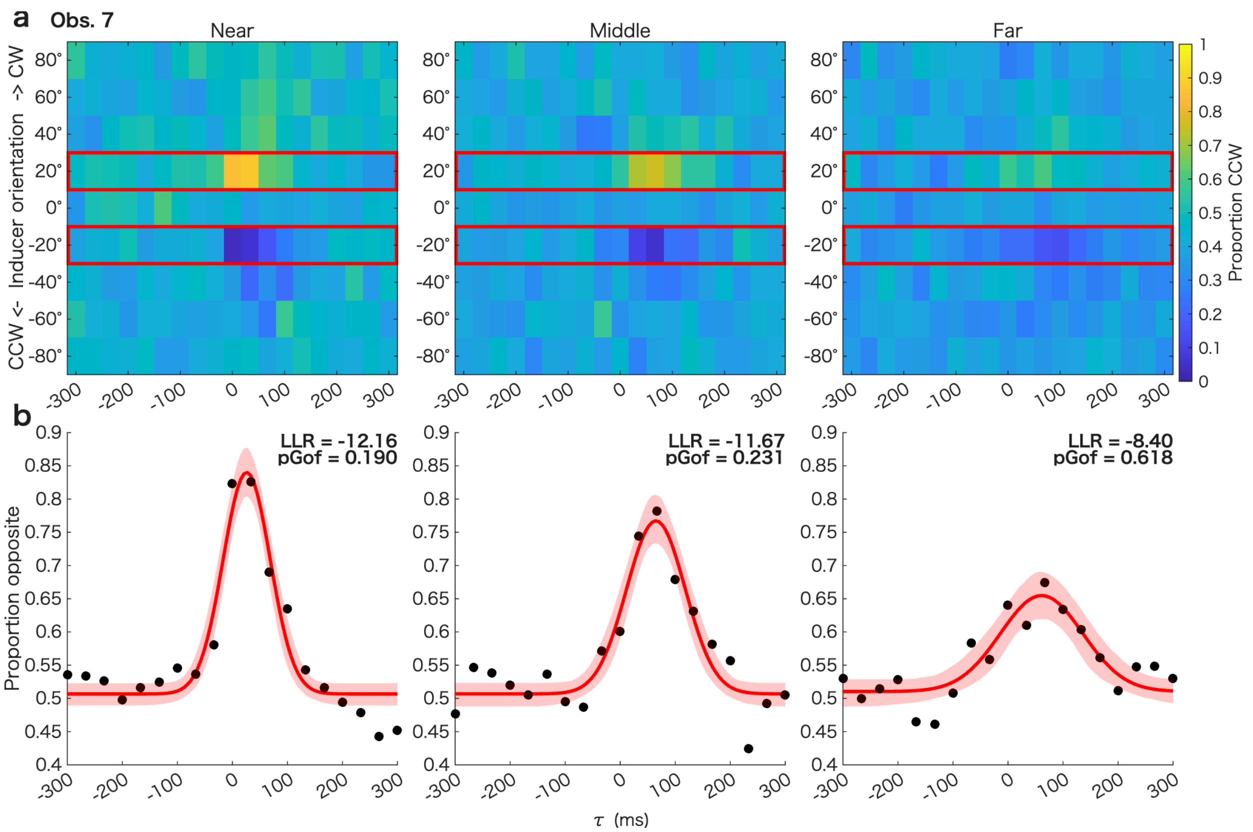
a
Obs. 7
Near
Middle
Far
CCW <- Inducer orientation -> CW
Proportion CCW
b
LLR = -12.16
pGof = 0.190
LLR = -11.67
pGof = 0.231
LLR = -8.40
pGof = 0.618
Proportion opposite
τ (ms)

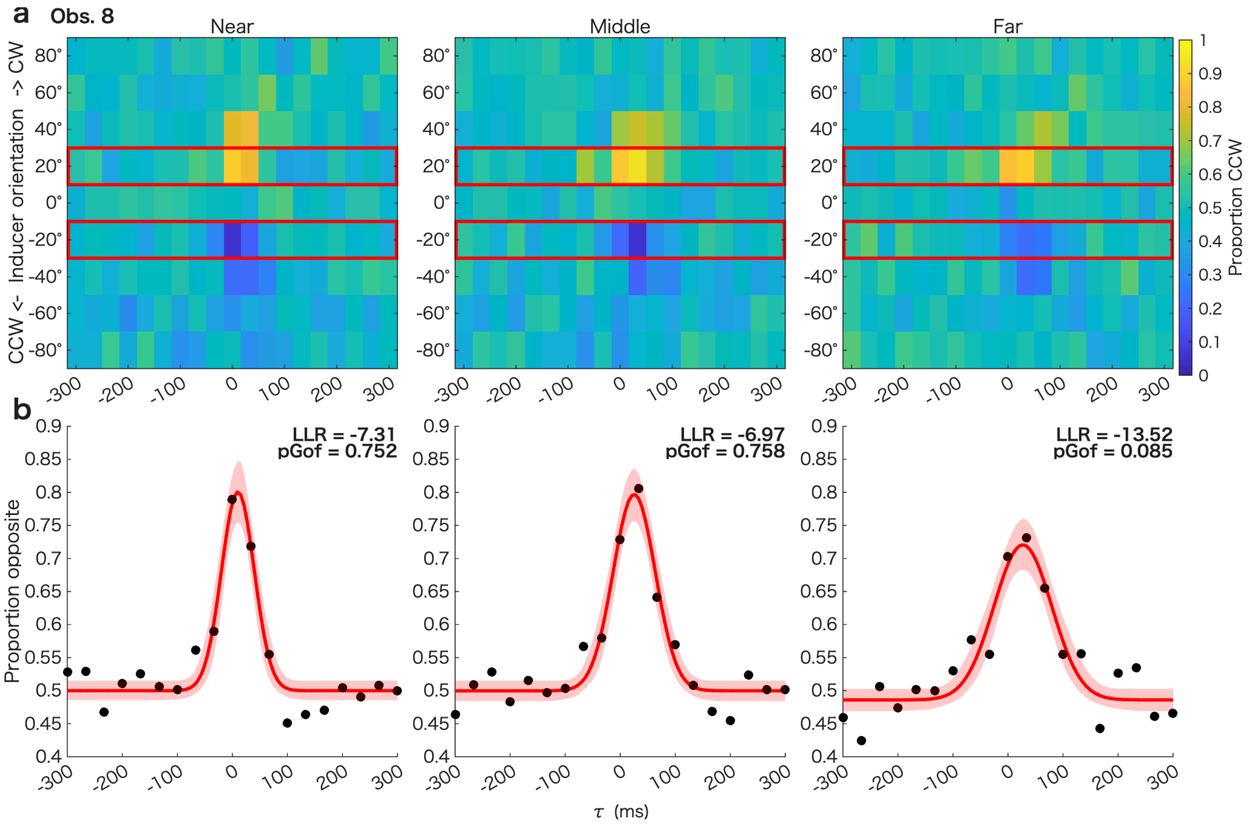
a
Obs. 8
Near
Middle
Far
CCW <- Inducer orientation -> CW
Proportion CCW
b
LLR = -7.31
pGof = 0.752
LLR = -6.97
pGof = 0.758
LLR = -13.52
pGof = 0.085
Proportion opposite
τ (ms)

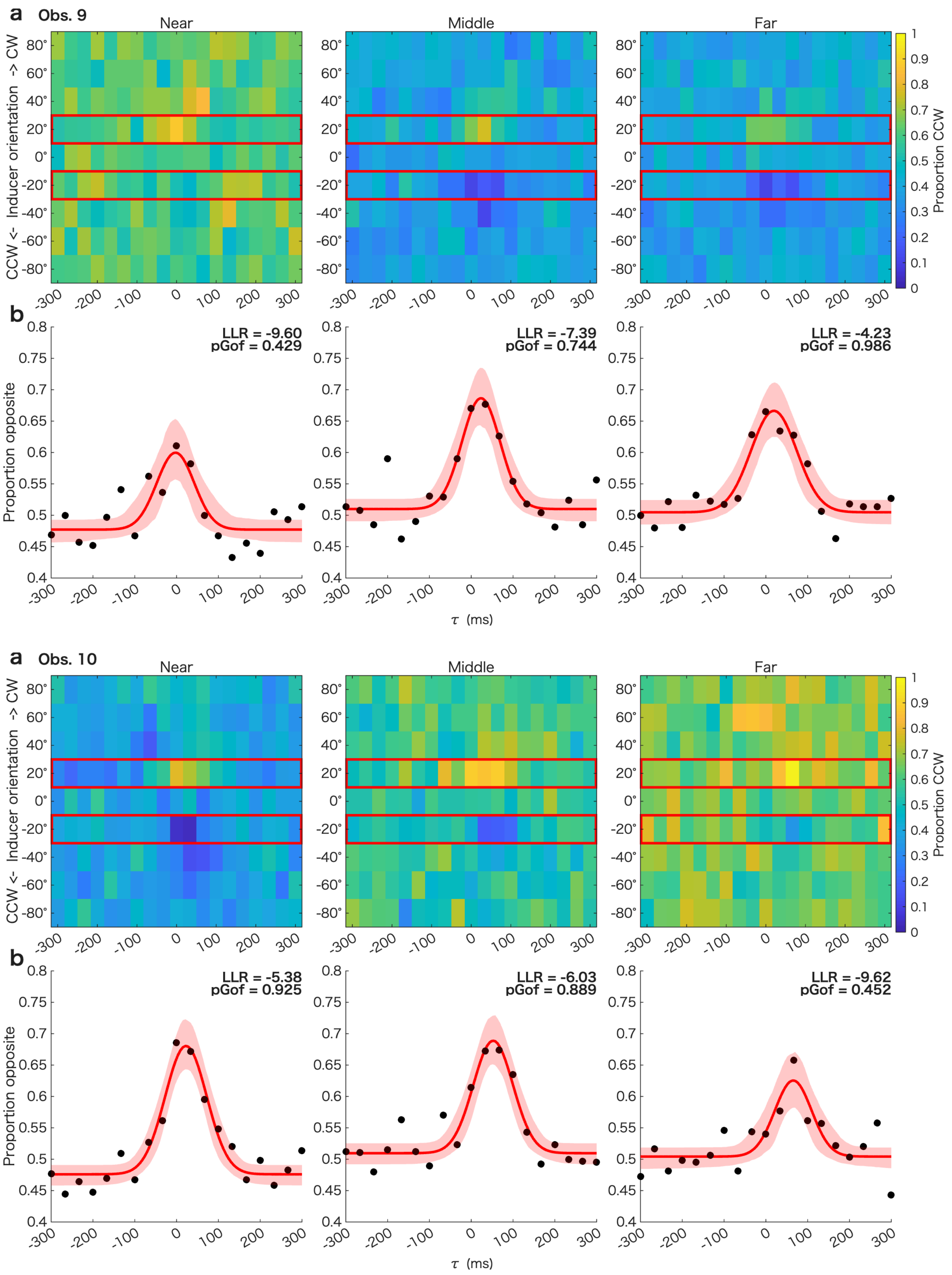


***Supplementary Fig. 1.*** *Correlograms and temporal windows of the tilt illusion in the near, middle, and far conditions, for each of ten observers. Conventions are the same as in Fig. 2.*

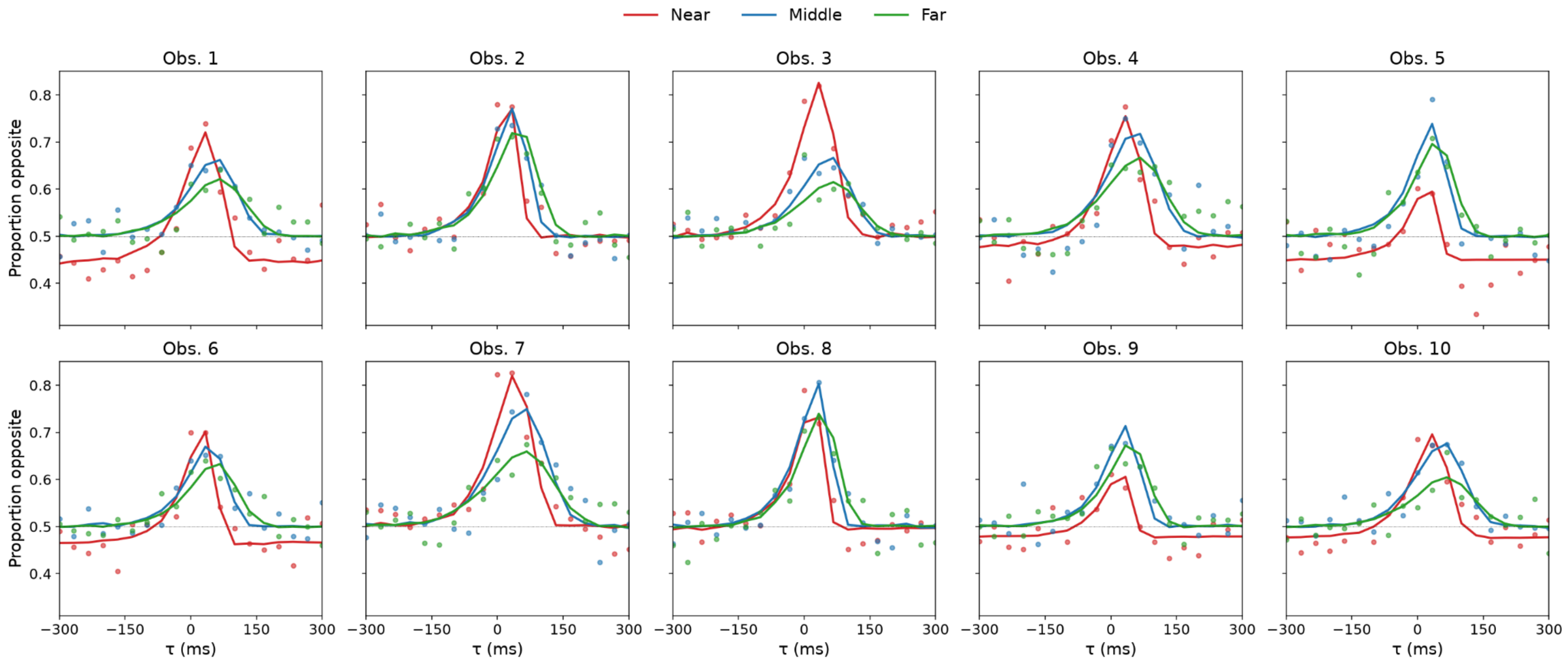


***Supplementary Fig. 2.*** *Predicted proportion opposite for ±20° inducers by the mechanistic active model. Each panel shows each observer's temporal windows: filled circles are the empirical proportion opposite and solid curves are the corresponding model predictions, computed as the expected proportion opposite over 40,000 simulated trials with randomly sampled inducer sequences. Distance is colour-coded (red, near = 2.0 dva; blue, middle = 3.5 dva; green, far = 5.0 dva).*

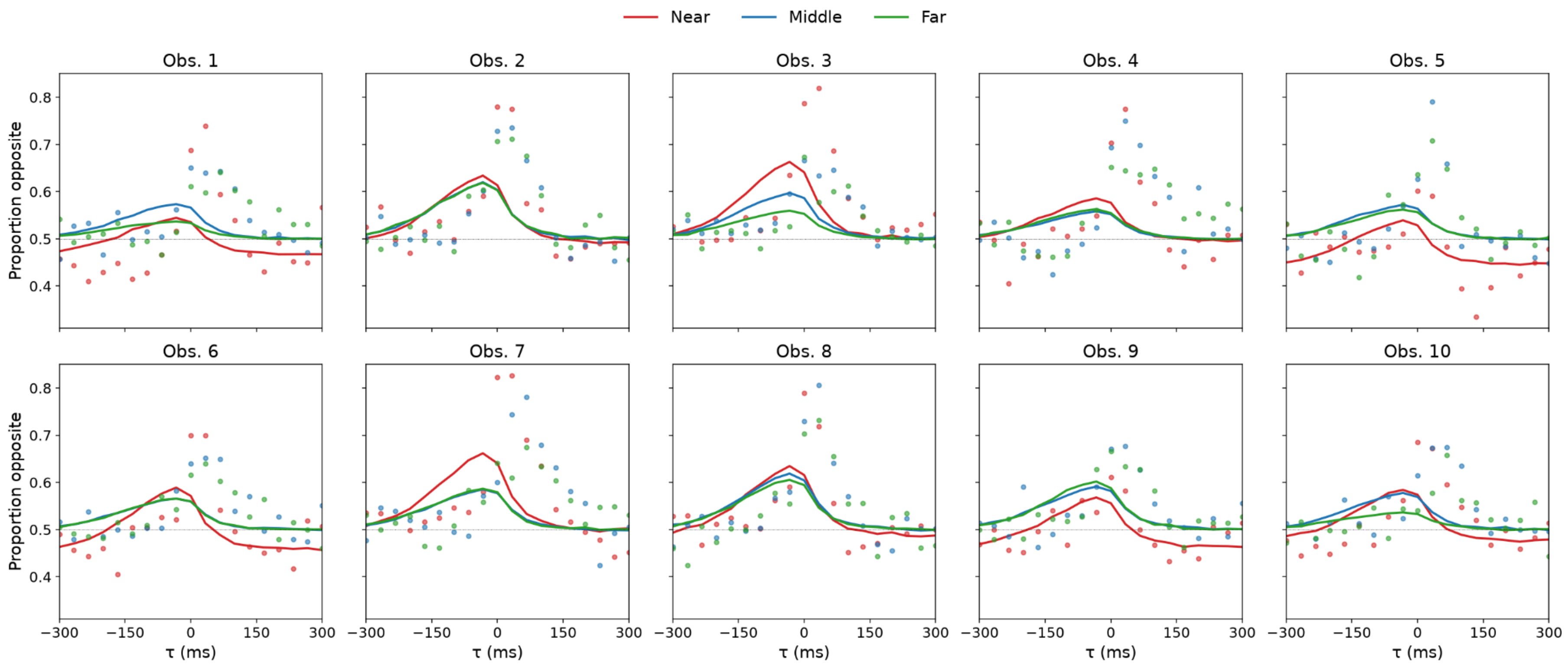


***Supplementary Fig. 3.*** *Predicted proportion opposite for ±20° inducers by the mechanistic passive model. Conventions are the same as in Supplementary Fig. 2.*

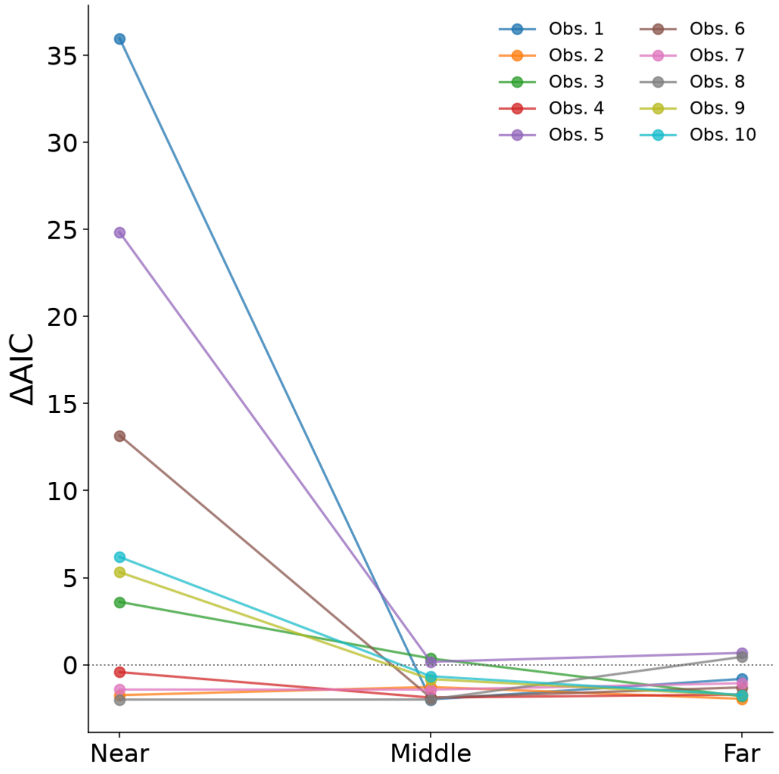


***Supplementary Fig. 4.*** *AIC difference between the three-parameter and four-parameter models. Positive ΔAIC values indicate higher AIC (i.e., poorer fit) for the three-parameter model compared to the four-parameter model. Different colours represent individual observers. In the near condition, AIC favoured the four-parameter model in the majority of observers.*

#### Supplementary Tables

**Supplementary Table 1.** Fitted parameters of the active model.

| Observer | $\alpha_{near}$ | $\alpha_{mid}$ | $\alpha_{far}$ | $v_h$ (m/s) | π |
|---|---|---|---|---|---|
| 1 | 15.37 | 5.35 | 6.85 | 0.073 | 0.283 |
| 2 | 5.91 | 6.90 | 6.91 | 0.127 | 0.008 |
| 3 | 9.65 | 5.68 | 6.70 | 0.071 | 0.000 |
| 4 | 8.91 | 8.47 | 10.61 | 0.072 | 0.113 |
| 5 | 3.74 | 5.39 | 5.28 | 0.141 | 0.261 |
| 6 | 6.46 | 4.21 | 4.23 | 0.103 | 0.180 |
| 7 | 10.68 | 13.22 | 14.43 | 0.063 | 0.000 |
| 8 | 5.09 | 7.38 | 6.39 | 0.157 | 0.028 |
| 9 | 2.62 | 4.66 | 4.49 | 0.138 | 0.117 |
| 10 | 6.07 | 6.25 | 5.95 | 0.071 | 0.120 |

Shared across observers: $\sigma_M = 0.359$; $k = 0.576$; $t_{reg} = 36.7$ ms.

**Supplementary Table 2.** Fitted parameters of the passive model.

| Observer | $\alpha_{near}$ | $\alpha_{mid}$ | $\alpha_{far}$ | $v_h$ (m/s) | π |
|---|---|---|---|---|---|
| 1 | 843.6 | 1063 | 2303 | 5.000 | 0.169 |
| 2 | 447.4 | 598.9 | 600.3 | 5.000 | 0.038 |
| 3 | 381.2 | 774.9 | 1372 | 5.000 | 0.000 |
| 4 | 842.2 | 1436 | 1298 | 5.000 | 0.017 |
| 5 | 527.5 | 1117 | 1293 | 5.000 | 0.287 |
| 6 | 362.2 | 1225 | 1220 | 5.000 | 0.219 |
| 7 | 384.4 | 911.2 | 891.6 | 5.000 | 0.000 |
| 8 | 396.5 | 604.2 | 683.4 | 5.000 | 0.074 |
| 9 | 521.7 | 829.7 | 743.2 | 5.000 | 0.196 |
| 10 | 560.6 | 1017 | 2337 | 5.000 | 0.124 |

Shared across observers: $\sigma_M = 0.442$; $k = 0.439$.
